# A cell-based platform for oxidative stress monitoring in motor neurons using genetically encoded biosensors of H_2_O_2_

**DOI:** 10.1101/2021.09.13.459724

**Authors:** Elizaveta I. Ustyantseva, Suren M. Zakian, Sergey P. Medvedev

## Abstract

**Background:** Oxidative stress plays an important role in the development of neurodegenerative diseases: it either can be the initiator or part of a pathological cascade leading to the neuron’s death. Although a lot of methods are known for oxidative stress study, most of them operate on non-native cellular substrates or interfere with the cell functioning. Genetically encoded (GE) biosensors of oxidative stress demonstrated their general functionality and overall safety in various live systems. However, there is still insufficient data regarding their use for research of disease-related phenotypes in relevant model systems, such as human cells.

**Methods:** We applied CRISPR/Cas9 genome editing to introduce mutations (c.272A>C and c.382G>C) in the associated with amyotrophic lateral sclerosis *SOD1* gene of induced pluripotent stem cells (iPSC) obtained from a healthy individual. Using CRISPR/Cas9, we modified these mutant iPSC lines, as well as the parental iPSC line, and a patient-specific *SOD1^D91A/D91A^* iPSC line with ratiometric GE biosensors of cytoplasmic (Cyto-roGFP2-Orp1) and mitochondrial (Mito-roGFP2-Orp1) H_2_O_2_. The biosensors sequences along with a specific transactivator for doxycycline-controllable expression were inserted in the “safe harbor” *AAVS1* (adeno-associated virus site 1) locus. We differentiated these transgenic iPSCs into motor neurons and investigated the functionality of the biosensors in such a system. We measured relative oxidation in the cultured motor neurons and its dependence on culture conditions, age, and genotype, as well as kinetics of H_2_O_2_ elimination in real-time.

**Results:** We developed a cell-based platform consisting of isogenic iPSC lines with different genotypes associated with amyotrophic lateral sclerosis. The iPSC lines were modified with GE biosensors of cytoplasmic and mitochondrial H_2_O_2_. We provide proof-of-principle data showing that this approach may be suitable for monitoring oxidative stress in cell models of various neurodegenerative diseases as the biosensors reflect the redox state of neurons.

**Conclusion:** We found that the GE biosensors inserted in the *AAVS1* locus remain functional in motor neurons and reflect pathological features of mutant motor neurons, although the readout largely depends on the severity of the mutation.

## INTRODUCTION

Redox reactions are part of the cellular metabolism. Normally, reactive oxygen species (ROS), emerging as a byproduct of such reactions, are quickly neutralized by antioxidant systems [1]. In oxidative stress, due to excessive production or disturbed utilization, ROS accumulate, subsequently leading to the cell malfunction [2]. An increasing number of the ROS molecules alter protein structure, change properties of membranes due to lipid peroxidation and cause DNA damage, which allows considering oxidative stress as one of the major mechanisms of degenerative disorders and aging [3, 4].

It is known that oxidative stress plays an important role in various pathologies, and its involvement in the development of neurodegenerative diseases is indisputable, although not always clear [5–7]. Oxidative stress takes a certain place in amyotrophic lateral sclerosis (ALS) – a disorder characterized by the inevitable death of motor neurons resulting in progressive paralysis [8]. The first ALS-associated gene, *SOD1*, has been discovered in 1993 [9]. *SOD1* encodes superoxide dismutase 1, the main component of the antioxidant system; thus, oxidative stress was proposed as the primary pathological mechanism of the disease [9]. Subsequent studies revealed that ALS has a much more complex etiology involving other genes and that only 10% of the cases are hereditary [10–12]. Moreover, it is considered now that not loss, but gain-of-function of mutant SOD1 underlies the ALS development [13]. Nonetheless, signs of oxidative damage have been found in both patients and model organisms, regardless of the initial cause of the disease, suggesting a universal role of redox imbalance in motor neuron damage [14–16].

Redox studies are often conducted by measuring key molecules such as glutathione, hydrogen peroxide, NADP^+^/NADPH, and others [17, 18]. Although dozens of chemical molecular probes have been developed for such analyses, most of them have low specificity, availability and interfere with the cellular processes [19, 20]. Genetically encoded (GE) biosensors are free from these flaws and can be applied for the same measurement as the molecular probes. GE biosensors are protein-based and delivered inside the cell in the form of nucleic acid. The cell produces biosensor molecules as long as the coding sequence is available. Since the biosensor molecules are produced inside the cell, they do not have the problem of availability, and therefore can be applied not only in cell cultures but also in more complex model systems, such as a whole animal or plant [21–23]. The nature of the genetically encoded biosensors allows easy modifications, e.g. addition of the tags that direct the biosensor to specific cellular compartments (the nucleus, mitochondria, endoplasmic reticulum, and plasma membrane) [24, 25]. Traditional methods for the GE biosensors research in cell culture apply their transient expression via plasmid delivery or viral-mediated integration of the biosensors sequences [4]. The first approach provides a high-intensity signal but does not allow prolonged experiments. The second one, on the contrary, provides a stable expression of the biosensor but does not guarantee reliable results since randomly integrated biosensors can disturb genome function. Furthermore, some may also speculate that a high level of the biosensor’s expression may alter cell functioning due to consumption of the target analytes [19]. Although many redox biosensors have been developed in recent years, only a few have been validated in model systems as suitable for studying disease-associated phenotypes [21,26–28].

Here, we developed a platform for monitoring oxidative stress in motor neurons. We used GE biosensors of cytoplasmic (Cyto-roGFP2-Orp1) and mitochondrial (Mito-roGFP2-Orp1) H_2_O_2_; a known marker molecule that reflects increased ROS production. The biosensors allow ratiometric measurement of hydrogen peroxide, and therefore relative oxidation in the corresponding compartments. The main advantage of the applied approach is that the biosensors’ sequences were inserted in the “safe harbor” *AAVS1* locus under the control of the doxycycline-dependent promoter, providing prolonged expression of the biosensors and preventing potential negative effect of off-target inserts. To investigate the functionality of the biosensors in these conditions, we generated isogenic induced pluripotent stem cell (iPSC) lines with two mutations affecting different parts of the ALS-associated *SOD1* gene. We found that the Cyto-roGFP2- Orp1 and Mito-roGFP2-Orp1 biosensors remain functional and reflect H_2_O_2_ levels in iPSC-derived motor neurons (Fig 1a). Moreover, we showed that a combination of G128R/K129X mutations, affecting exon 5 of *SOD1*, results in rapid accumulation of H_2_O_2_ and an aberrant response to exogenous H_2_O_2_ in mature motor neurons, but it was not observed for the D91A mutation.

**Fig. 1.**
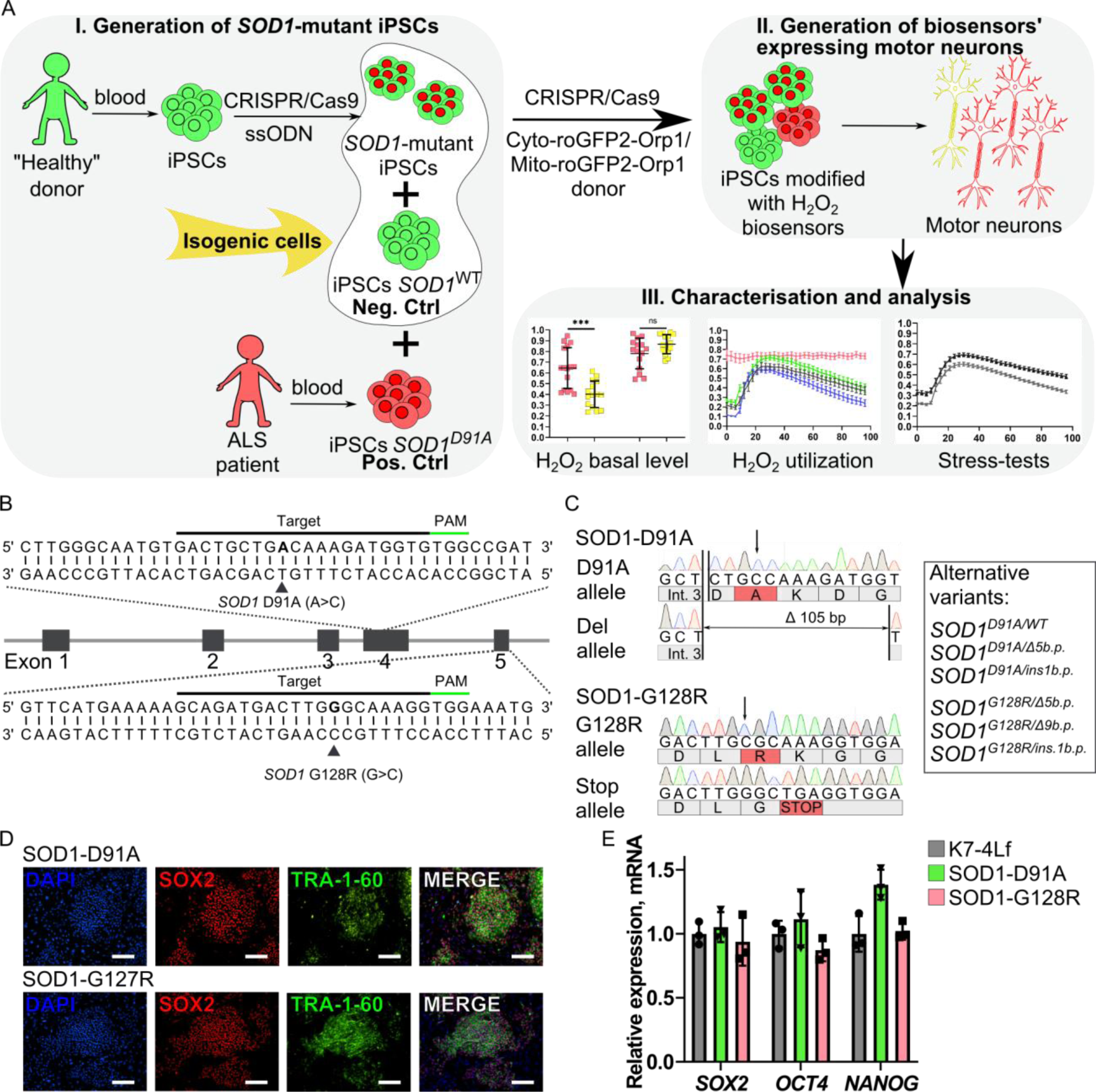
Generation of isogenic *SOD1*-mutant iPSC lines. **A** Schematic representation of the experimental design. iPSCs – induced pluripotent stem cells; Neg. Ctrl – negative control; Pos. Ctrl – positive control; ssODN – single- stranded oligodeoxynucleotide. **B** Schematic of the *SOD1* gene with partial sequences of the exons 4 and 5. Protospacers designed for CRISPR/Cas9-mediated double-stranded breaks are underlined with black lines, PAM – with green lines; target mutations are in bold and marked with black triangles. **C** Partial *SOD1* sequences of the exons 4 and 5 of the SOD1-D91A and SOD1-G128R iPSC lines. Int. 3 – intron 3. In the box: list of clones with an alternative *SOD1* variants obtained in the study. **D** Immunocytochemistry of the SOD1-D91A and SOD1-G128R iPSC lines. The cells are positively stained for pluripotency markers: transcriptional factor SOX2 and surface antigen TRA-1-60. Nuclei are visualized with DAPI, scale bar 100 μm. **E** RT-qPCR analysis of mRNA expression of *SOX2*, *OCT4*, and *NANOG* in the SOD1-D91A and SOD1-G128R iPSCs. Data (N = 3) are normalized to the parental iPSC line (K7-4Lf) and presented as the mean ± standard deviation. Characterization of the SOD1-D91A and SOD1-G128R iPSC lines in detail is in the Supplementary Fig. S2.

## METHODS

### IPS cell culture

IPSCs were maintained onto mitotically inactivated mouse embryonic fibroblasts in KnockOut DMEM (Gibco) with 15% knockout serum replacement (Gibco), 0.1 mM non-essential amino acids (Gibco), penicillin/streptomycin (Lonza), 1 mM GlutaMAX-I, and 10 ng/mL bFGF at 37°C and 5% CO_2_. IPSC cells were dissociated with TrypLE (Gibco) and split at 1:10 twice a week in the iPSC medium supplemented with 10 ng/ml Y-27632. Original iPSC lines were derived from the patient (iALS) with a diagnosed hereditary form of ALS [29] and a healthy individual (K7-4Lf) who had no associations with any genetic disease [30] **(Supplementary Table 1)**.

### CRISPR/Cas9 design and generation of *SOD1* mutant iPSC lines

The guide RNAs (gRNAs) targeting sequences in exons 4 and 5 of the *SOD1* gene and the *AAVS1* locus were designed using the web-based tool http://crispr.mit.edu [31], with a selection of gRNAs with high-quality scores to avoid possible off-targets **(Supplementary Table 2)**. We used CRISPR/Cas9 ribonucleoprotein (RNP) complexes to induce double-strand breaks in the target sites. The Alt-R® crisprRNA and tracrRNA were obtained from IDT (Integrated DNA technologies), and Cas9 protein was expressed in *E. coli* and purified according to the previously published protocol [32]. The RNP complexes were assembled according to the manufacturer’s instructions before the cell transfection. For the introduction of the c.272A>C and c.382G>C mutations we used appropriate SOD1 RNP (20 pmol tracrRNA + 20 pmol crRNA (SOD1- 4/SOD1-5) + 20 pmol Cas9) complexes mixed with 100 pmol of D91A ssODN (single-stranded oligodeoxynucleotide) donor or G128R ssODN donor **(Supplementary Table 2)**. The cells were passed 24 hours before the transfection in the iPSC medium supplemented with Y-27632 (10 ng/ml). On the day of transfection, the cells were dissociated with TrypLE, strained through a 70 μm cell strainer (Miltenyi biotec), counted, centrifuged at 200g for 5 min, and resuspended in R buffer (Neon Transfection System, Invitrogen) according to the manufacturer’s instructions. 10 ul of the suspension was taken to the electroporation by Neon Transfection System with the following impulse settings: 1100 V, 30 ms, 1 pulse. The cells were seeded onto feeder-coated 4 cm^2^ dishes in the iPSC medium supplemented with Y-27632 (10 ng/ml). The next day, the cells were dissociated with TrypLE, strained through the cell strainer, and subcloned on three 96-well plates three cells per well for propagation and analysis. Genomic DNA of the survived clones was obtained and analyzed for the presence of the target mutations.

### Screening of iPSC clones for the c.272A>C (D91A) and c.382G>C (G128R) substitutions

To detect the c.272A>C mutation, we designed primers for tetra-primer ARMS (amplification-refractory mutation system) PCR screening using http://primer1.soton.ac.uk/primer1.html **(Supplementary Table 2**) [33] and performed touchdown 3-step PCR: annealing at 68-64 °C for 9 cycles, then at 64 °C for 21 cycles. The PCR products were analyzed in 2% agarose gel. Clones positive for mutant allele presence were further examined by Sanger sequencing **(Supplementary Table 2)**. To detect the c.382G>C mutation, we designed a pair of primers that amplify the target locus of the *SOD1* gene and two fluorescent probes targeting either wild-type or mutant sequence **(Supplementary Table 2)**. Using LightCycler 480 (Roche), we analyzed the clones and selected those who had strong signals from the mutant-targeted probe. The target mutation was further confirmed by Sanger sequencing **(Supplementary Table 2)**. The clones used in the experiments were characterized according to the Human Pluripotent Stem Cell Registry standards with the protocols described earlier [34].

### Generation, selection, and screening of iPSC clones with target *AAVS1* inserts

To insert Cyto-roGFP2-Orp1, Mito-roGFP2-Orp1 and transactivator in *AAVS1*, we used AAVS1 RNP (100 pmol tracrRNA + 100 pmol AAVS1 crRNA + 100 pmol Cas9) mixed with 5 μg of donor plasmids mix, containing equimolar amounts of transactivator donor (pAAVS1-Neo- M2rtTA, Addgene # 60843) + pCyto-roGFP2-Orp1-donor or pMito-roGFP2-Orp1-donor. The cells were prepared as it was described earlier and resuspended in R buffer. We mixed 100 μl of the cell suspension, RNP complexes and donor plasmids and performed transfection using Neon Transfection System (2× reaction per experiment). The cells were then seeded onto feeder-coated 10 cm^2^ dishes in the iPSC medium supplemented with Y-27632 (10 ng/ml) and maintained until small colonies formed, for 2-3 days prior to the selection. For the selection of subclones with the target biosensor and transactivator inserts, we supplemented the iPSC medium with puromycin dihydrochloride (Sigma-Aldrich) for 3 days. Then we replaced the antibiotic with neomycin sulfate (Sigma-Aldrich) and incubated the cells for 4-5 more days. Antibiotics concentrations were determined for each cell line by titration before the experiment. At the end of the selection, we added doxycycline hyclate (2 μg/ml, Sigma-Aldrich) and examined the remained clones for the presence of fluorescent signal from the biosensors’ roGFP2 (reduction-oxidation sensitive green fluorescent protein 2) using the Nikon Eclipse Ti2-E (Nikon) microscope. The clones positive for the roGFP2 expression that survived double antibiotic selection were manually harvested into separate dishes for maintaining and analysis. We extracted genomic DNA from these iPSC clones and analyzed for the presence of the target and off-target inserts of the donor plasmids using PCR with specific primers **(Supplementary Table 2)**.

### Immunocytochemistry

Immunocytochemistry was performed on iPSCs and motor neurons (ChAT – differentiation day 20 and 28; ISL1 and MNX1 – differentiation day 28). The cells were fixed in 4% formaldehyde solution (Sigma-Aldrich) for 10 min at room temperature (RT), permeabilized with 0.5% Triton X-100 (Sigma-Aldrich) for 30 min at RT, and then incubated with blocking buffer (1% bovine serum albumin (BSA) in PBS, Sigma) for 30 min at RT. After, the cells were incubated with specific primary antibodies overnight at 4 °C. The appropriate secondary antibodies were added for 1.5-2 h incubation at RT. All antibodies were diluted in blocking buffer, and the cell nuclei were visualized with DAPI (1 μg/ml solution in PBS; Sigma-Aldrich). The antibodies and their dilution ratios are listed in the **Supplementary Table 3**. Micrographs were captured using either Nikon eclipse Ti-E microscope (Nikon) and NIS Elements software or LSM-780 (Zeiss) microscope and ZEN black software.

### Motor neuron differentiation

The iPSCs were seeded onto dishes coated with Matrigel-ESQ (Corning) in E8 (Gibco) medium and maintained in feeder-free conditions for at least 2 passages prior to differentiation. Motor neuron differentiation was performed according to the previously published 4-step protocol [35]. For neural patterning, the E8 medium was changed to basal neuronal differentiation medium (NDM): F12/DMEM:Neurobasal – 50:50, 0.5× N2 supplement, 0.5× B-27 supplement, 2 mM GlutaMAX, and 0.1 mM ascorbic acid. At the first step of differentiation, the NDM was supplemented with 2 μM CHIR99021 (StemRD), 2 μM SB431242 (Selleckchem), and 2 μM DMH1 (Tocris) for 6 days. Then the cells were dissociated with Accutase (Gibco) and plated 1:6 onto Matrigel-coated surface in the NDM medium, supplemented with CHIR99021, SB431242, DMH1, 0.5 μM Purmorphamine (Stegment), and 0.1 μM retinoic acid (Sigma) for another 6 days (the second step). After, the cells were dissociated with Accutase and cultured in the suspension on agarose-coated (non-adherent) dishes in the NDM medium, supplemented with 0.1 μM Puromorphamin and 0.5 μM retinoic acid for 6 days (the third step). Then the cells were dissociated with Accutase to a single-cell suspension, strained through the 70 μm cell strainer, and seeded in the NDM medium, supplemented with 0.1 μM Puromorphamin, 0.5 μM retinoic acid, and 0.15 μM Compound E (EMDMillipore) either on Matrigel-coated surfaces or inside a layer of the Matrigel for 10-day maturation (fourth step). Depending on the experiment, the cells were seeded in different settings. For flow cytometry and mRNA expression analysis, we seeded 1.5×10^5^ cells/cm^2^ onto Matrigel-coated 60 mm Petri dishes; for immunocytochemistry – 5×10^4^ cells/cm^2^ – onto Matrigel-coated cell imaging coverglasses (Eppendorf); for axon measurement – 1.5×10^4^ cells/cm^2^ – onto Matrigel-coated cell imaging coverglasses. After dissociation between each step, cells were transferred in the NDM medium, supplemented with Y-27632 (10 ng/ml).

The medium was replaced every day. The cells were supplemented with doxycycline (2 μg/ml) every other day during the differentiation unless otherwise indicated.

For starvation induction, we used Neuronal Deficit Medium (NDefM, F12/DMEM:Neurobasal – 50:50, 1x N2 supplement, 2 μg/ml doxycycline hyclate). For antioxidant-deprivation assay and excitotoxicity induction assay, MN were incubated in the neuronal maintenance medium without antioxidants (F/D:Neurobasal – 50:50, 0.5× N2 supplement, 0.5× B-27 Supplement without antioxidants), supplemented with 0.5 μM retinoic acid, and 0.15 μM Compound E.

### Preparation of live motor neuron samples for microscopy

Motor neurons have a low surface attachment, which makes prolonged microscopy experiments difficult. Therefore, in the biosensors experiments, the cells were seeded for maturation on the cell imaging coverglasses inside a layer of 33% Matrigel. To do so, we resuspended the cells in the 1.5× cold fourth step medium (supplemented with 15 ng/ml Y-27632), making a suspension with 1.5×10^5^ cells/75 μl (1.5×10^5^ cells/well). Then, we mixed 75 μl of the cell suspension with 35 μl of growth factor reduced Matrigel (Corning) and quickly applied 100 μl of the mix onto the surface of the chilled cell imaging coverglass, standing on a cold tube cooling rack turned upside down and covered with a paper towel. Using the tip of a pipette, we carefully spread the mix over the surface and left it on the cooling rack for 10 minutes to let the cells fall to bottom before the Matrigel polymerized. After, we transferred the coverglasses on top of the working surface and left them for another 10 minutes, and, then, the coverglasses were carried over to a CO_2_ incubator for 1h for Matrigel stabilization. After 1 hour, we added 300-400 μl of the fourth step medium supplemented with Y-27632 (10 ng/ml) on top of the stabilized Matrigel layer **(Supplementary Fig. S1)**.

### Reverse-transcription quantitative PCR (RT-qPCR)

Total RNA was extracted from iPSC and motor neurons with Trizol reagent (Invitrogen). The reverse transcription of 1 μg of total RNA was performed with 5x RT-buffer mix with M-MuLV- RH reverse transcriptase (Biolabmix) and random hexamer primer (Invitrogen) and diluted 1:10 in MilliQ H_2_O. To measure the endogenous gene expression of pluripotency factors, motor neurons’ markers, and transgene expression the qPCR analysis was performed using LightCycler 480 (Roche). Gene expression of the iPSC markers in the *SOD1* mutant iPSCs was normalized to the *B2M* housekeeping gene and compared to the original iPSC line using the ΔΔCt-method. *ISL1*, *CHAT,* and *MNX1* genes expression in motor neurons was normalized to the mean of the *GAPDH*, *HPRT1,* and *RPL13* housekeeping genes and compared to expression in iPSC sample using the ΔΔCt-method. The roGFP2 and rtTA transgenes expression in motor neurons and iPSCs was normalized to the mean of *GAPDH*, *HPRT1*, and *RPL13* housekeeping genes and compared to the expression in motor neuron sample that was not treated with doxycycline during differentiation. The primers used are listed in the **Supplementary Table 2**.

### Flow cytometry analysis

To identify the proportion of motor neurons in the differentiation, we dissociated the cells with Accutase on the day 20 of the differentiation protocol, resuspended in cold PBS, and centrifuged at 400 g for 5 minutes (the same settings were used for all subsequent centrifugation steps). The pellet was resuspended in 1 ml cold 4% formaldehyde solution and incubated on ice for 10-15 minutes. After, we added 1 ml cold PBS, centrifuged the cells, discarded supernatant, resuspended the pellet in 1 ml ice-cold 100% methanol, and incubated it for 10-15 minutes on ice. Then, the pellet was washed twice with flow cytometry staining buffer (1% BSA, 0.2 μM EDTA, in PBS) and resuspended in it to 1×10^6^ cells/ml concentration. 100 μl of the cell suspension was incubated with anti-ISL primary antibodies overnight at 4 °C. The cells were washed with the flow cytometry staining buffer and incubated with the secondary antibodies for 30 minutes at RT. Cells were analyzed using FACSAria (BD Biosciences). Unlabeled cells and isotype-labeled cells were used as controls.

### Fluorescence intensity measurement

To measure the biosensors’ fluorescence intensity, we obtained images of the MN using a Zeiss LSM-780 confocal laser scanning microscope (Pan-Apochromat 20× objective) adjusted for visualization of a green dye: excitation with 488 nm argon laser, emission collection at 500-530 nm. Using the ImageJ software, we measured the mean intensity of fluorescence on each image and calculated corrected total cell fluorescence (CTCF) with the formula: CTCF = Integrated Density – (Area occupied by cells × Mean fluorescence of background readings) to reduce the background contribution. The results were obtained from six separate experiments, which are represented by the MN derived from different iPSC clones.

### Axon measurement

The immature motor neurons were seeded on the cell imaging coverglasses in a low density (1.5×10^4^ cells/cm^2^) and grown for 2 days in the fourth step medium supplemented with neurotrophic factor (NTF) cocktail: IGF1 (PeproTech, 10 ng/ml), CNTF (PeproTech, 10 ng/ml), BDNF (PeproTech, 10 ng/ml). Doxycycline (2 μg/ml) was added to induce the roGFP2 expression. RoGFP2 served as a label, marking the cellular contour and processes of the live neurons. We obtained mages for each cell line with the Nikon Eclipse Ti-2E microscope (20× objective, FITC channel). Using ImageJ, we manually measured the length of the longest processes (axons) of free-lying neurons with visible ends. Only axons with the length more than twice the size of the neurons’ bodies were considered for the analysis. If the neuron had two long processes, the longest one was considered for the measurement. The mean length of the axons was calculated based on the data obtained from the differentiation of three separate iPSC clones for each genotype present.

### Image acquisition

The general procedure for the redox biosensors measurement was described earlier by B. Morgan and colleagues [36]. Although, we modified the protocol to be more suitable for cultured motor neurons.

#### Cells and solutions preparation

To measure H_2_O_2_ utilization in real-time, we replaced the medium in the neurons with the neuronal deficit medium the day before the experiment, unless otherwise indicated. 1 hour before the experiment the medium has been removed from the analyzed wells (so as not to disturb the Matrigel layer with neurons inside), and replaced with warm HBSS + Ca^2+^, Mg^2+^. The cells were incubated in a CO_2_ incubator for removal of the residual medium components from the Matrigel layer. Then, the old HBSS has been almost completely removed (with residual of ∼ 50 μl/well) and replaced with the fresh HBSS (∼ 300 μl/well).

The stock and working solutions were prepared freshly on the day of the experiment. The water stock solutions of 1 M DTT (Sigma-Aldrich) and 0.2 M diamide (Sigma-Aldrich) were made from a powder; 10 μM H_2_O_2_ stock solution was made from hydrogen peroxide solution (30% w/w, Sigma-Aldrich).

From these stock solutions, the working solutions has been prepared:

1. 10 μl l 1M DTT + 190 μl HBSS;
2. 5 μl 0.2M diamide +195 μl HBSS;
3. 2 μl 10 μM H_2_O_2_ + 198 μl HBSS.

#### Microscopy settings

We used a confocal Zeiss LSM-780 laser scanning microscope (Pan-Apochromat 20× objective) equipped with the 488 nm argon laser, the 405 nm UV diode laser, and a climate chamber connected with the temperature and CO_2_ control modules. Cell imaging coverglass with the cells was placed on the stage without a lid, covered with a CO_2_ cover, and left for 5-10 minutes for temperature equilibration. Tubes with the working solutions were also put inside the climate chamber. During the equilibration, the cells were visualized with transmitted light to achieve a stable focus. Actual microscopy settings varied between the samples and required customization for obtaining a quality image. Table 1 describes the initial microscopy setup. An image did not contain oversaturated pixels and had a high noise-to-signal ratio.

**Table 1.** Initial microscopy setup (Zeiss LSM-780) for imaging of Cyto-roGFP2-Orp1 and Mito- roGFP2-Orp1 expressing motor neurons

| Mode | Channel mode |
| --- | --- |
| *Channels:* |  |
| Switch track every | «Frame» |
| Track 1 | 405 nm (DAPI, roGFP2ox) + T-PMT (Transmission) |
| Track 2 | 488 nm (EGFP, roGFP2red) |
| *Light Path:* |  |
| Track 1+ Track 2 | 500-530 nm |
| *Acquisition mode:* |  |
| Scan mode | Frame |
| Frame size | 512x512 (1024x1024 for photo) |
| Scanning speed (pixel dwell) | 9 (1,58 μsec) |
| Averaging | 1 |
| Bit depth | 16 bit |
| Zoom | 0.6 |
| *Channel/Laser settings:* |  |
| 405-nm line |  |
| Attenuator (transmission) | 5% |
| Pinhole | 74.5 |
| Gain | 800 |
| Digital gain | 1 |
| 488-nm line |  |
| Attenuator (transmission) | 2% |
| Pinhole | 74.5 |
| Gain | 800 – Cyto-roGFP2-Orp1, 700 - Mito-roGFP2-Orp1 |
| Digital gain | 1 |

#### Biosensor calibration using DTT/diamide

To determine the states of the maximum oxidation and reduction possible for the biosensors, we treated the cells with diamide and DTT, respectively, using the following procedure:

1. In the mode “Position” set up three fields of view in each of the two adjacent wells and saved their coordinates;
2. Set the microscope to the “time series” mode with 18 cycles of 90 s;
3. Started the time series experiment and paused it after cycle 3; added 50 μl of the warm DTT working solution into one well (final concentration 5 μM) and 50 μl of the warm diamide working solution into the other well (final concentration 0.5 μM), trying not to disturb the coverglass and the cell layer; resumed the time series;
4. Waited for the cells to become fully reduced/oxidized (usually 8-10 minutes): the signal in each sample reached the plateau. In the DTT well intensity of the 488 nm signal slightly increased, while intensity of the 405 nm signal slightly decreased; in the diamide well intensity of the 488 nm signal intensity considerably decreased, while intensity of the 405 nm signal considerably increased.

The calibration procedure has been performed for every sample before the other experiments, and the values of the maximum oxidation/reduction were used for the dynamic range calculation and data normalization.

#### The basal H_2_O_2_ level measurement and the measurement of H_2_O_2_ utilization in real-time

For the measurement of the Cyto-roGFP2-Orp1 and Mito-roGFP2-Orp1 signals in motor neurons, we applied the microscope settings determined during the calibration. For the measurement of basal H_2_O_2_ levels in cytoplasm in mitochondria, we obtained images from MN derived from the K7-4Lf, SOD1-D91A, SOD1-G128R, and iALS iPSC lines. Starvation was induced by changing the medium to the neuronal deficit medium. Rescue of the SOD1-G128R MN was performed by the addition of a neurotrophic factor (NTF) cocktail: IGF1 (10 ng/ml), CNTF (10 ng/ml), and BDNF (10 ng/ml) to the culture medium. To measure H_2_O_2_ utilization in real-time, we applied the same approach as for calibration, but with 33 cycles, each 3 minutes long. We prepared the cells as described, mounted the coverglasses on the stage, and equilibrated the temperature in the climate chamber. We added 50 μl of the warm working H_2_O_2_ solution (final concentration 10 μM) to the cells after cycle 2-3, and then continued the time series for another 30 cycles (1.5 hours).

### Excitotoxicity induction assay

To induce glutamate-mediated excitotoxicity, we incubated MN in the neuronal maintenance medium supplemented with 0.5 μM Retinoic acid, 0.15 μM Compound E, 20 μM monosodium glutamate (Sigma-Aldrich), and 100 μM l-trans-pyrrolidine-2,4-dicarboxylic acid (PDC, Sigma-Aldrich) for 5 days, changing the medium every other day. After 5 days of incubation, we obtained images of the treated MN and non-treated control MN. The data obtained at the end of the experiment were normalized to the starting oxidation values, measured before the glutamate addition, to describe the changes that emerged during the experiment.

### Data normalization

The images were saved as 16-bit .tiff files and processed by ImageJ. For the analysis, the images were converted to the 32-bit format. Single pictures were split to the 405 nm (roGFP2_ox_), 488 nm (roGFP2_red_), and transmitted light channels. The intensity of the 405 and 488 channels was thresholded, and values below the threshold were set to “not a number” (NaN). A ratio image was created by dividing the 405 nm image to the 488 nm image (roGFP2_ox_/roGFP2_red_), and the mean intensity of the resulting image was measured. The time series images were processed similarly. The files were imported in ImageJ as stacks, converted to the 32-bit format, split using the “Stacks-Shuffling-Deinterleave” plugin. The threshold was adjusted for both channels, and the values below the threshold were set to “not a number” (NaN). A resulting ratio images were created by dividing the 405 nm image to the 488 nm and measured. The roGFP2_ox_/roGFP2_red_ ratios of the single images were used to describe the basal level of H_2_O_2_. The roGFP2_ox_/roGFP2_red_ ratios obtained from the time series were used for measurement of H_2_O_2_ utilization in real-time. Data obtained from different lines were normalized to a fraction of the maximum oxidation/reduction state according to the formula (1):

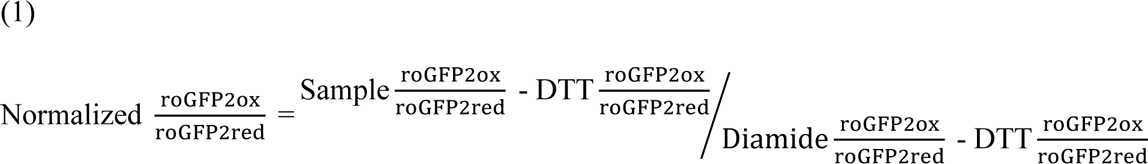

The normalized data were used for visualization, comparison and statistical analysis.

### Maximum oxidation and recovery rate calculation

The maximum biosensor’s oxidation was calculated by subtraction of the initial normalized value of the roGFP2ox/roGFP2red ratio recorded at “0” time point from the normalized maximum oxidation value of the roGFP2ox/roGFP2red ratio obtained after H_2_O_2_ addition (usually after 30 minutes of recording). The recovery rate was calculated by subtraction of the final normalized value of the roGFP2ox/roGFP2red ratio recorded at the end of the time series from the normalized maximum oxidation value with subsequent division of the resulting numbers to the time interval between these two time points (in hours).

### Statistics

The data derived from at least three different clones of the same genotype were collected for the comparisons; except for measurements of H_2_O_2_ utilization in starvation assay and Mito-roGFP2- Orp1 reaction to H_2_O_2_, where data were collected from the MN derived from one iPSC clone. Analysis of the axon length was performed using one-way ANOVA with the Bonferroni correction for multiple comparisons. Analyses of mRNA expression, fluorescence intensity and all the information obtained from the biosensors were performed using the Mann-Whitney U-test with the Bonferroni correction for multiple comparisons where necessary. Averaged data are represented in the form of scatter plots, or line graphs as the mean ± standard error of the mean (the mean ± S.E.M) or as the mean ± standard deviation (the mean ± S.D.). The number of experimental replicates is denoted as N. Statistical significance is defined as * p < 0.05, ** p < 0.01, *** (or ###) p < 0.001, **** (or ####) p < 0.0001.

## RESULTS

### Introduction of the *SOD1* D91A and G128R mutations in IPSCs of the clinically healthy donor

It is known that *SOD1* has more than 140 mutations associated with ALS, and they define clinical features of the disease such as its manifestation age, rate of symptoms progression, presence of additional neurological symptoms, etc [37]. Several theoretical studies [38, 39] hypothesized that severity of the symptoms is highly dependent on the position of a mutation in the sequence and its effect on the protein folding and stability. We designed two CRISPR/Cas9 guide RNA targeting the sequences in the exons 4 and 5 of *SOD1* and two ssODN donor templates necessary for the introduction of c.272A>C and c.382G>C single nucleotide mutations that lead to the D91A and G128R substitutions, respectively, in the SOD1 polypeptide (Fig. 1b, **Supplementary Table 2)**. The c.272A>C mutation is known for its relatively mild character with late manifestation and long progression, while c.382G>C is characterized by extremely rapid development [40, 41].

We introduced these mutations using CRISPR/Cas9, into a well-characterized control iPSC line (K7-4Lf), obtained earlier from a clinically healthy individual [30] **(Supplementary Table 1)** and recovered 66 clones for D91A variant and 124 clones for G128R variant. We screened those clones by tetra-primer ARMS PCR (for D91A) or qPCR with fluorescent probes (for G128R) and found 6 (9.1%) and 4 (3.2%) clones, respectively, presumably positive for the target mutations. The target mutations were further confirmed by Sanger sequencing. As a result, a number of clones with different *SOD1* allelic variants were obtained (Fig. 1c). Since no homozygous clones were found, we chose clones with *SOD1^D91A/del105^* (SOD1-D91A) and *SOD1^G128R/K129*^* (SOD1-G128R) variants for subsequent experiments. It was suggested that the destruction of one of the alleles will create a more severe phenotype and make biosensors’ measurements more robust. These iPSC lines demonstrated features specific for pluripotent cells: they expressed specific genes and proteins (OCT4, NANOG, SOX2, and TRA1-60), positively stained for alkaline phosphatase, and were able to generate three germ layer derivatives. The cells also retained a normal 46, XX karyotype and were free of mycoplasma contamination (Fig. 1d-e, **Supplementary Fig. S2)**.

### Generation of transgenic IPSC lines with the inserts of genetically encoded biosensors of H_2_O_2_ via CRISPR/Cas9-mediated *AAVS1* targeting

We introduced the sequences of two genetically encoded biosensors: Cyto-roGFP2-Orp1 and Mito-roGFP2-Orp1, measuring H_2_O_2_ levels in the cytoplasm and mitochondria, respectively, in the genome of SOD1-D91A, SOD1-G128R, and the isogenic control line (K7-4Lf) to obtain stable expression. Additionally, we introduced the same sequences in the genome of a patient-specific iPSC line (iALS) previously generated in our lab from the patient, diagnosed with a hereditary form of ALS with homozygous D91A mutation in *SOD1* [34]. This cell line does not share the same genetic background with the *SOD1-*mutant and control iPSC lines but serves as an external positive control.

The Tet-On expression system applied for the biosensors’ expression consists of two elements [42]: the biosensor’s sequence that follows the tetracycline-dependent promoter and the specific transactivator (rtTA, reverse tetracycline-controlled transactivator) essential for the controlled expression of the target genes. To deliver these elements in the cell’s genome, we used biallelic target insertion in the safe harbor *AAVS1* locus (the first intron of the *PPP1R12C* gene) via CRISPR/Cas9 (Fig. 2a). The donor plasmid containing the rtTA, homologous arms, and neomycin resistant gene for the selection was obtained from the vendor (Addgene) [43]. The biosensors’ donor plasmids were designed and constructed in the lab [29]. These donor plasmids contained the tetracycline-dependent promoter, followed by the Cyto-roGFP2-Orp1 and Mito- roGFP2-Orp1 sequences, as well as ∼800 base long homology arms, and puromycin resistance gene for the selection of the target clones (**Supplementary Fig. S3).** We delivered both donor plasmids along with the CRISPR/Cas9 RNPs in the IPSCs and selected them using appropriate media supplemented with neomycin and puromycin. Among the survived clones, we manually picked up and subcloned those who positively responded to the doxycycline (tetracycline derivative) addition with the biosensors expression and screened the clones for the presence of target biallelic insertion in the *AAVS1* locus by PCR (Fig. 2b-c). Further, clones positive for the target insertions were screened for additional off-target donor integration by PCR using specific pairs of primers that only detect non-integrating parts of the donor plasmids, implying that the presence of the PCR-product indicates an off-target insertion. We obtained from 3 to 17 separate iPSC clones positive for the target and negative for the off-target insertions for each cell line (Fig. 2d). All transgenic iPSC clones used further in the experiments were stained positive for the specific pluripotent cell markers SOX2 and SSEA4 (**Supplementary Fig. 4)**.

**Fig. 2.**
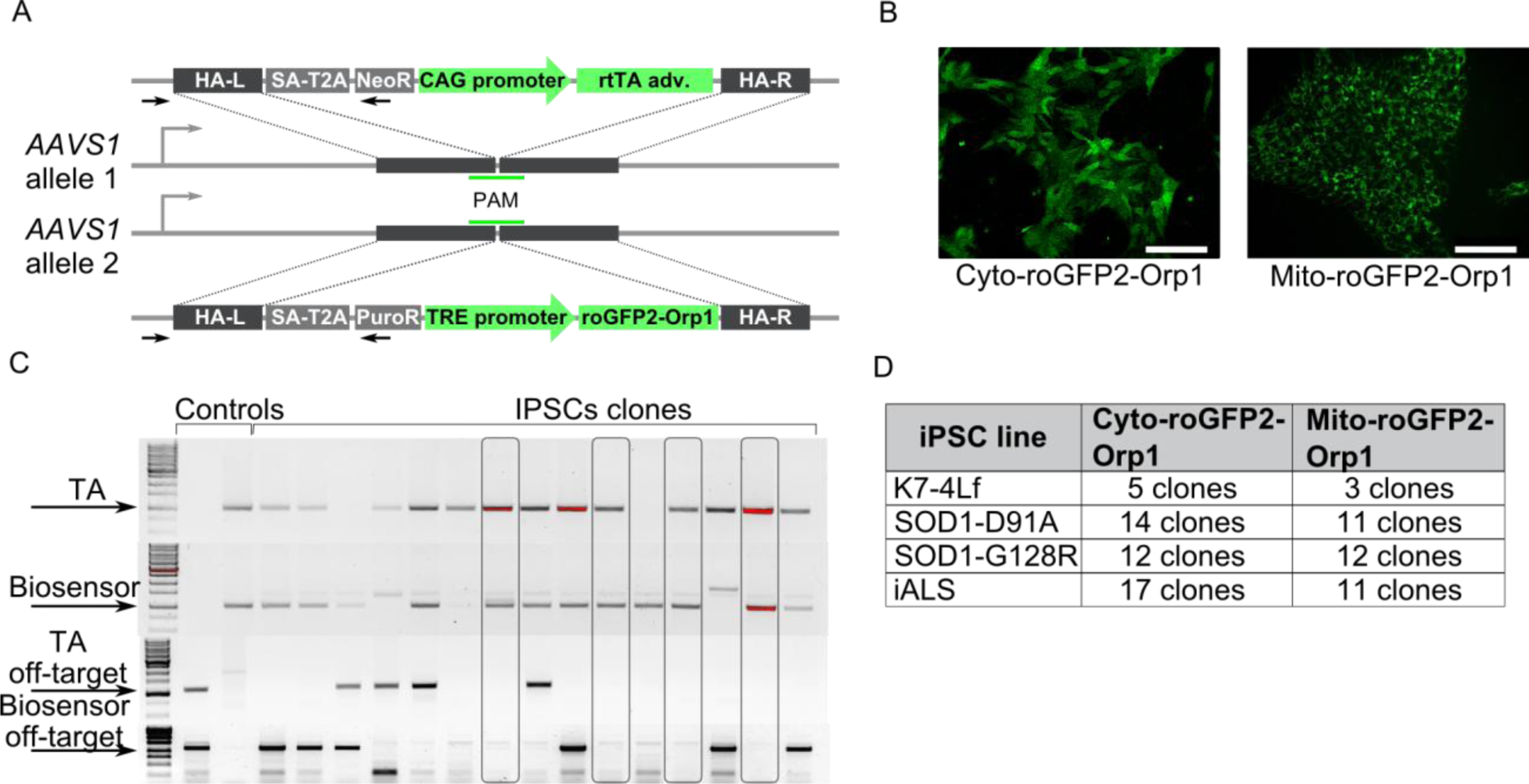
Generation of iPSCs expressing the Cyto-roGFP2-Orp1 and Mito-roGFP2-Orp1 biosensors. **A** Schematic of the biosensor and transactivator for doxycycline-controllable expression inserts in the *AAVS1* locus. In one allele, the homologous arms (HA-L – left, HA-R – right) flank the splice acceptor site (SA), T2A peptide, the neomycin-resistance gene (in the open reading frame of *PPP1R12C*), and rtTA under control of the CAG promoter. In another allele, the homologous arms flank the splice acceptor site (SA), T2A peptide, the puromycin-resistance gene (in the open reading frame of *PPP1R12C*), and the biosensor’s sequence under control of the TRE-promoter. The primers used to detect the target insertions (one – inside the insert, another – outside the left homologous arm, in the *AAVS1* locus) are represented as black arrows. **B** Representative images of live iPSCs expressing the Cyto-roGFP2-Orp1 (left) or Mito-roGFP2-Orp1 (right) biosensors 24 h after the addition of 2 mg/ml doxycycline in the medium. Scale bar 100 μm. **C** Representative Image of the iPSC clones analyzed for the target (two top gels) and off-target (two bottom gels) inserts. The arrows mark target PCR products. The clones positive for the target and negative for the off-target inserts are in frames. **D** List of iPSC clones obtained for each type of the biosensor and *SOD1* variant.

### Motor neurons derived from the IPSCs with mutant *SOD1* show impaired axon growth

We utilized a previously described protocol of highly efficient motor neuron differentiation (Fig. 3a) [35] that allowed us to obtain iPSC-derived motor neurons with a characteristic morphology within 30 days. For each type of iPSC (K7-4Lf, iALS, SOD1-D91A, and SOD1-G128R), we differentiated three separate clones. All motor neurons derived from the iPSC were positively stained for ChAT, ISL1, and MNX1 and expressed mRNA of these proteins on comparable levels (Fig. 3b-c, **Supplementary Fig. S5**). We analyzed MN differentiation efficiency on the day 20 of differentiation by counting the ISL^+^ cells using flow cytometry. The efficiency ranged between 89 and 95% with the 91.9±7% ISL^+^ cells for K7-4Lf, 91.1±2% for SOD1-D91A, 89.3±3.8% for SOD1-G128R, and 94.7±1.5% for iALS (Fig. 3d).

**Fig. 3.**
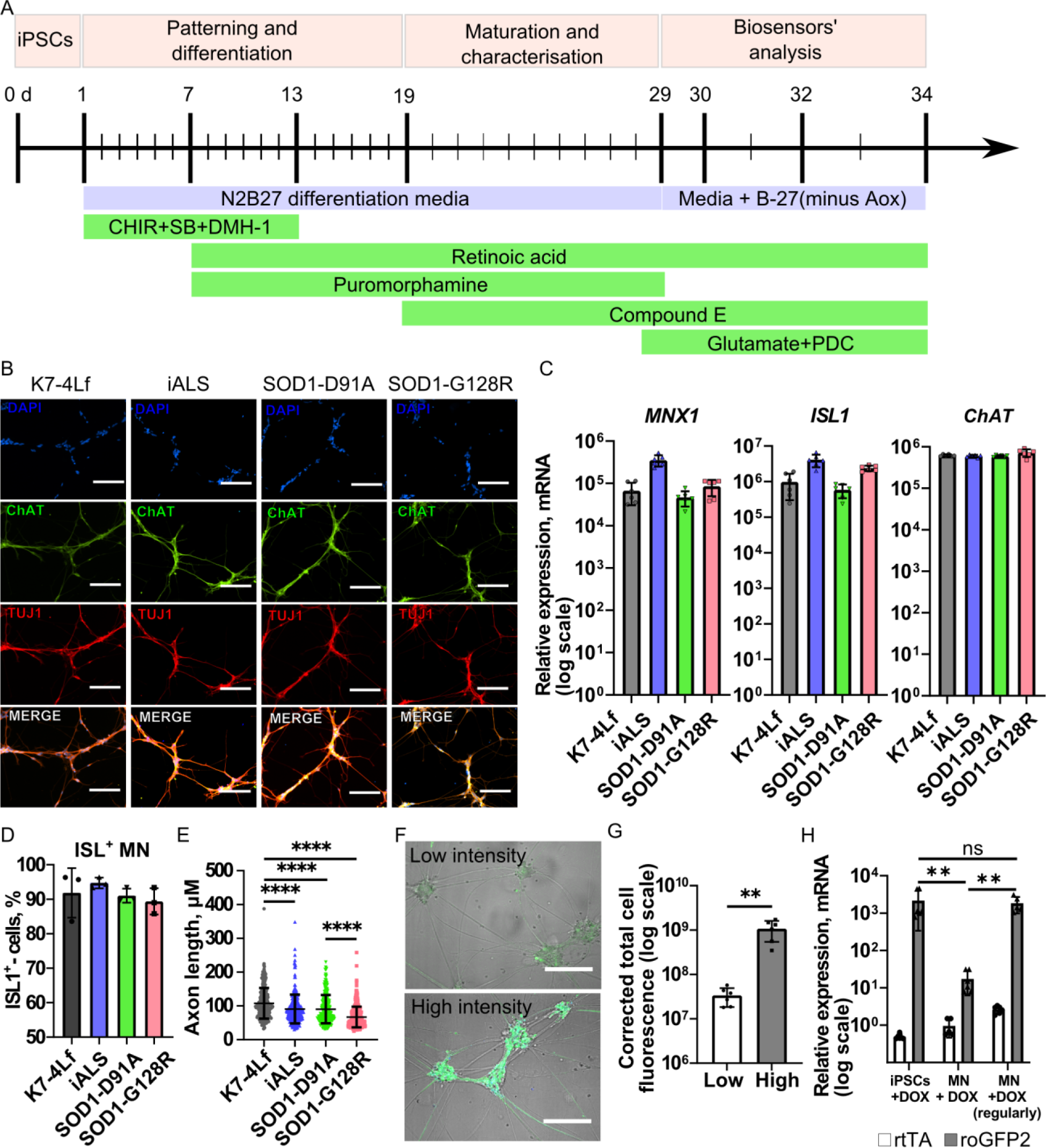
Generation of iPSC-derived motor neurons expressing the Cyto-roGFP2-Orp1 and Mito-roGFP2-Orp1 biosensors. **A** Schematic of the differentiation protocol and a timeline for MN characterization and analysis. The main stages of the protocol are denoted above the timeline. The medium composition (basal medium and small- molecule inhibitors) for every stage is indicated below the timeline. **B** Representative Images of the MN positively stained for the general neuronal marker β-III-Tubulin (TUJ1) and mature MN marker ChAT on the differentiation day 20. Nuclei are visualized with DAPI, scale bar 100 μm. **C** RT-qPCR analysis of mRNA expression of *MNX1*, *ISL1*, and *ChAT* in K7-4Lf, iALS, SOD1-D91A, and SOD1-G128R derived MN on the differentiation day 29. Data (N = 6) are normalized to the expression in the K7-4Lf iPSC line and presented as the mean ± standard deviation (S.D.). **D** Quantification of flow cytometry analysis of ISL^+^ cells performed on the differentiation day 20. Data (N = 3) are the mean ± S. D. **E** Axon length quantification (differentiation day 21). Data (N = 257, 270, 409, 397 for K7- 4Lf, iALS, SOD1-D91A, and SOD1-G128R, respectively) are presented as the mean ± S.D. **F** Representative Images of the MN lost (top) and retained (bottom) the Cyto-roGFP2-Orp1 expression during the differentiation. Scale bar 100 μm. **G** Quantification of the Cyto-roGFP2-Orp1 fluorescence intensity in MN lost and retained the Cyto-roGFP2-Orp1 expression. Data (N 6) are the mean ± S.D. **H** RT-qPCR analysis of the transactivator (rtTA) and biosensors (roGFP2) mRNA expression 48 h after 2 μg/ml doxycycline addition. MN + DOX – MN differentiated in the absence of doxycycline, MN + DOX (regularly) – MN differentiated with regular doxycycline supplementation, iPSCs +DOX – iPSCs expressing biosensor. Data (N 6) are the mean ± S.D. ** p < 0.01, **** p < 0.0001 with the Bonferroni correction for multiple comparisons where necessary.

To characterize MN obtained, we measured axonal length of the derived MN on the day 21 of differentiation in low-density culture. The mean length of the axons in SOD1-D91A and iALS MN was 90.7±42.6 μm and 90.4±41.9 μm, respectively, which was significantly lower than in control K7-4Lf MN (107.8±45.5 μm). The mean axon length in SOD1-G128R (67±30.6 μm) was even lower than in SOD1-D91A and almost forty percent lower compared to the control K7-4Lf. This suggests the presence of the pathological effect of the mutations introduced in *SOD1*, as well as a different level of severity of these mutations (Fig. 3e).

### The biosensors’ expression declines during differentiation

Although the *AAVS1* site is located in the intron of a transcriptionally active gene and was described previously as suitable for stable expression of transgenes [44], we have discovered that the motor neurons did not always retain detectable fluorescence level of the biosensors at the terminal stages of the differentiation, and this did not depend on the particular cell line used for the differentiation (Fig. 3f-g). Analysis of the expression level of rtTA and roGFP2 in the MN that lost the biosensor’s signal revealed that the terminally differentiated MN expressed mRNA of the rtTA at the same level as the corresponding IPSCs from which they were obtained, while the expression of the biosensor’s roGFP2 was decreased by two orders, suggesting that the biosensor’s promoter was selectively inhibited (Fig. 3h). It is known that the differentiation process is accompanied by chromatin remodeling [45]. Since the rtTA expression is constitutive, while biosensors require tetracycline-derivatives to activate transcription, we suggested that the active state of the promoter during differentiation prevents its inhibition. We have been supplementing the differentiation medium with doxycycline every other day from the first day of differentiation to keep the biosensor’s promoter in an active state. This resulted in a more stable expression of the biosensors’ mRNA on a level comparable to the IPSC (Fig. 3h) as well as in a high level of the fluorescence intensity.

### Target insertion of the genetically encoded biosensors of H_2_O_2_ in the *AAVS1* locus does not affect their basic properties

The main principle of the ratiometric H_2_O_2_ biosensors is based on the roGFP2 ability to change its fluorescent properties. Two cysteine residues positioned close to the roGFP2 chromophore form a disulfide bond under oxidation, leading to structural changes that influence the protein fluorescence [46]. Thus, the reduced roGFP2 (roGFP2red) have excitation maximum at 490 nm, the oxidized (roGFP2ox) – at 400 nm, with the emission maximum at 510 nm for both forms. It is possible to obtain a signal from each form separately by exciting the biosensor at two wavelengths; the intensity of these signals, in turn, allows estimating ***relative*** oxidation of the roGFP2, i.e. the proportion of the oxidized molecules in the cytoplasm (for Cyto-roGFP2-Orp1) or mitochondria (for Mito-roGFP2-Orp1). Fusion of the roGFP2 to the thiol peroxidase Orp1 creates a redox relay in which H_2_O_2_ specifically oxidizes Orp1, and then this state is passed to the roGFP2, with subsequent Orp1 reduction (Fig. 4a) [46]. Since the reaction is reversible, the biosensors reflect not only the steady-state level of H_2_O_2_ in the cell but also dynamic changes in H_2_O_2_ caused by certain events.

**Fig. 4.**
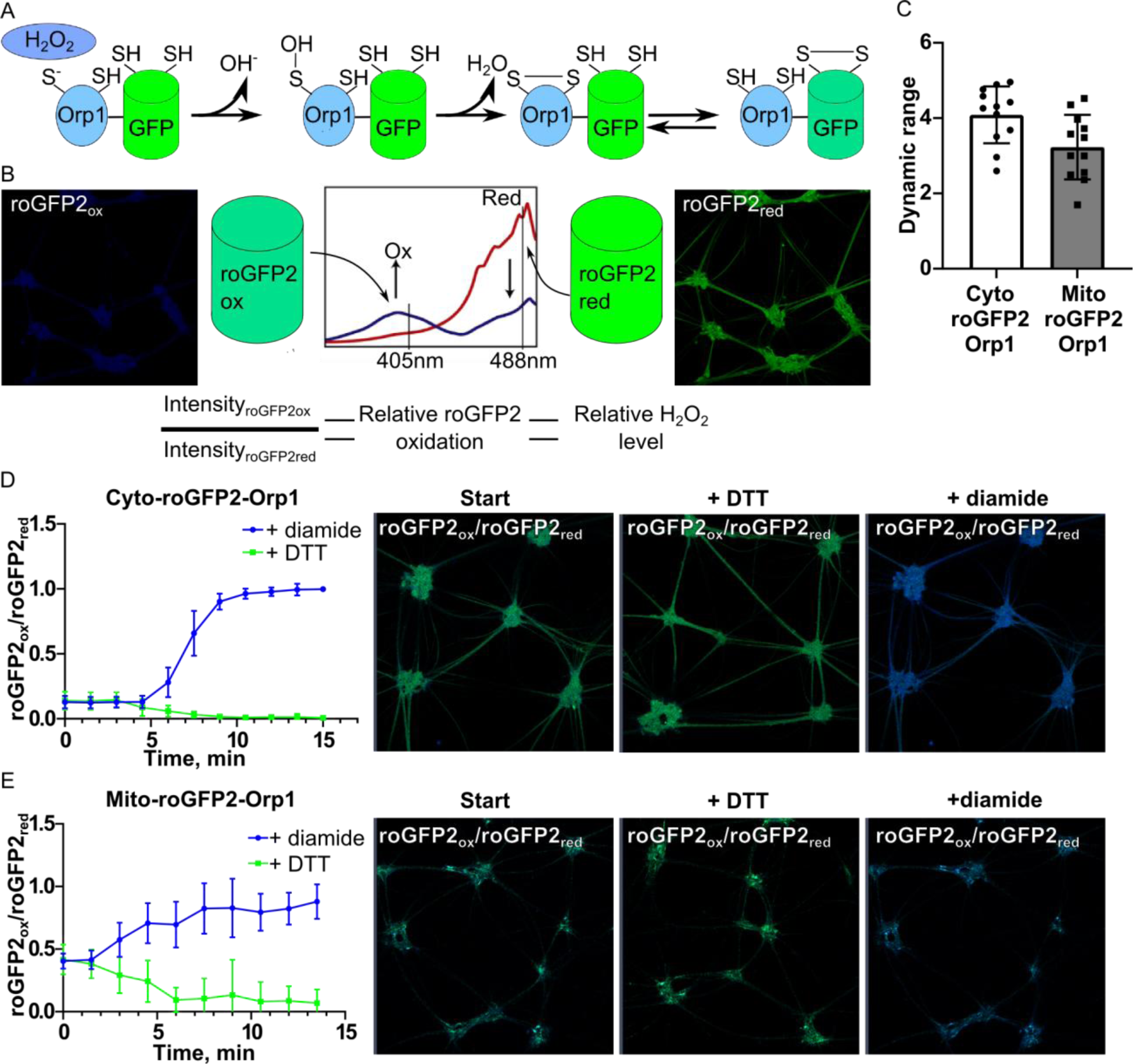
Evaluation of the Cyto-roGFP2-Orp1 and Mito-roGFP2-Orp1 functionality in iPSC-derived motor neurons. **A** Interaction of roGFP2-Orp1 with H_2_O_2_ (According to Morgan et al. [36]). The Image demonstrates how H_2_O_2_ triggers oxidation of the roGFP2 via Orp1 thiol peroxidase. **B** Fluorescence spectra of roGFP2 (top) in the fully reduced (roGFP2_red_) or fully oxidized (roGFP2_ox_) states with the representative images of MN obtained by excitation at the 405 nm (blue pseudocolor) and 488 nm (green pseudocolor) wavelengths; the equation for calculation of the relative biosensor’s oxidation (bottom). **C** Dynamic range of the Cyto-roGFP2-Orp1 and Mito-roGFP2-Orp1 biosensors expressed in MN. Data (N 12) are the mean ± standard deviation (S.D.). The reaction of the Cyto- roGFP2-Orp1-expressing (**D**) and Mito-roGFP2-Orp1-expressing (**E**) MN to the diamide and DTT addition with the representative images of the intact MN (Start), fully reduced (+ DTT) and fully oxidized (+ diamide) MN. Data (N = 9) are normalized to the values of the maximum oxidation and reduction and presented as the mean ± S.D. The Images present an overlap of the 405 nm (oxidized roGFP2) and 488 nm (reduced roGFP2) channels colored in blue and green pseudocolors, respectively.

For the ratiometric H_2_O_2_ biosensors, dynamic range – the difference between the fully oxidized and fully reduced biosensor – reflects the scale of the signals that can be distinguished in the experiment (Fig. 4b), and according to the literature, this number varies from 3 to 8 [23,36,47]. We measured the dynamic range of the Cyto-roGFP2-Orp1 and Mito-roGFP2-Orp1 biosensors in every MN sample used in the work (Fig. 4c). To do that, we added to the cells either oxidizing agent diamide or reducing agent DTT. Afterward we recorded changes in the biosensors’ signal in real-time. The biosensors quickly reacted to the addition of the oxidizing and reducing agents, reaching the plateau 8-10 minutes after the chemicals were added (Fig. 4d-e). The calculated dynamic range was 4±0.33 for Cyto-roGFP2-Orp1 and 3.3±0.61 for Mito-roGFP2-Orp1 and stayed within acceptable values.

### B-27 supplement deprivation affects mitochondrial level of H_2_O_2_ regardless of the genotype

The basal level of H_2_O_2_ reflects redox balance and general condition of the cell. To obtain information about the redox state of the MN, we performed live imaging on the differentiation day 29. We did not observe any differences between K7-4Lf and SOD1-D91A MN: relative oxidation level of the cytoplasm and mitochondria was the same for both types of neurons (Fig. 5a). No differences were also detected between these two cellular compartments. The cytoplasmic H_2_O_2_ level in iALS MN was slightly lower compared to the control (K7-4Lf), but the mitochondrial H_2_O_2_ level was the same. We suggested that these findings are probably due to the different MN origin of iALS MN since the SOD1-D91A oxidation values did not differ from the control, although this statement requires additional research. SOD1-G128R MN, however, demonstrated a 2.7 times higher level of H_2_O_2_ in the cytoplasm and a 5 times higher level of the mitochondrial H_2_O_2_ compared to the control. Together with an overall higher level of the oxidation in both these compartments, we observed that the mitochondrial level of H_2_O_2_ was twice as high as the cytoplasmic. Notably, that the cytoplasmic oxidation measured at the differentiation day 20 (immature MN) was the same for all neurons, suggesting that the SOD1- G128R MN dysfunction, probably caused by the *SOD1* mutation, occurs only in mature MN (Fig. 5b). To correct the observed phenotype, we added a combination of neurotrophic factors (NTF) to the culture medium during SOD1-G128R MN maturation (differentiation days 19-29). This resulted in significant decrease in cytoplasmic oxidation to the normal level. The mitochondrial level of H_2_O_2_, however, was not affected by the NTF addition (Fig. 5c).

**Fig. 5.**
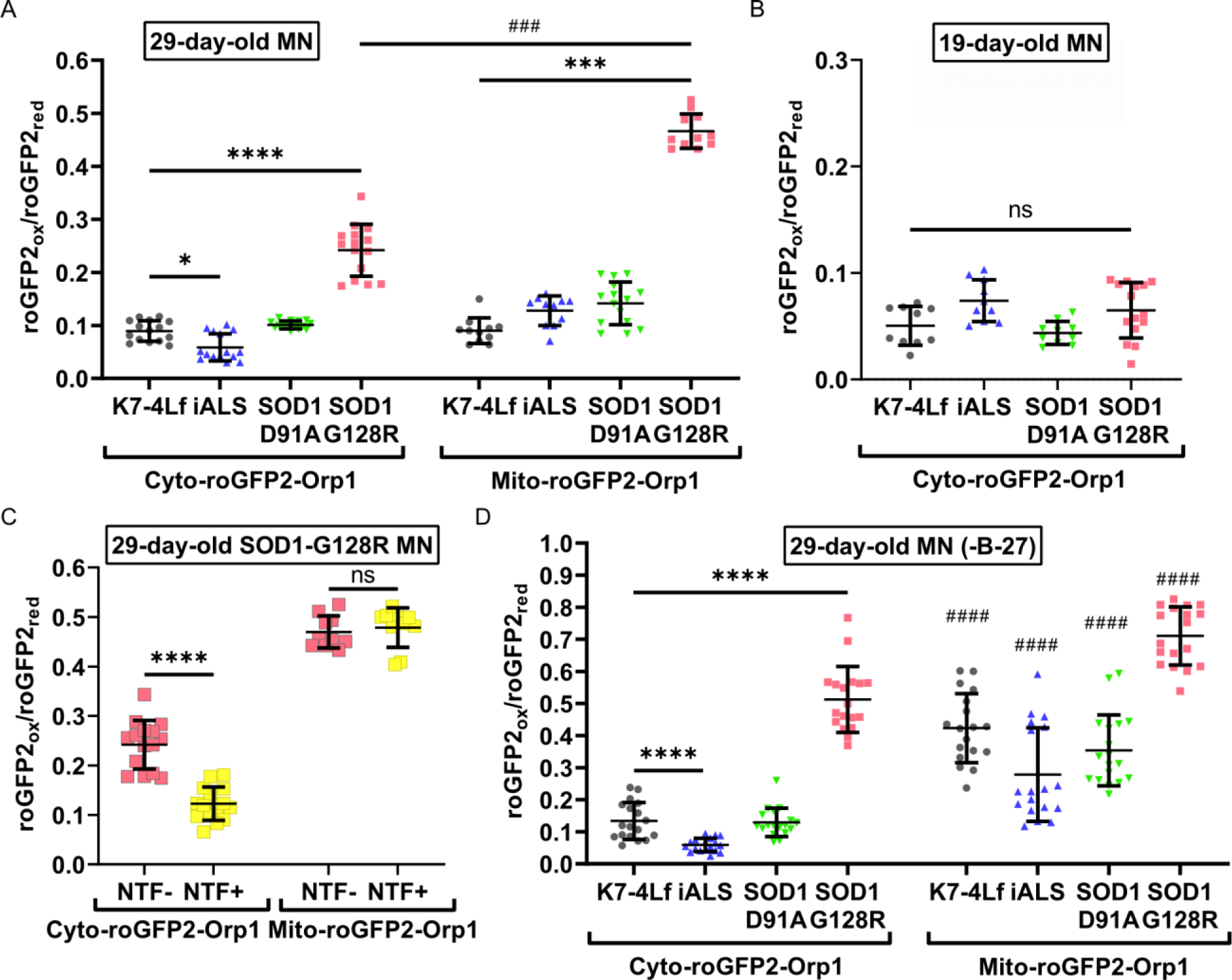
Basal levels of H_2_O_2_ in the cytoplasm and mitochondria of the Cyto-roGFP2-Orp1 and Mito-roGFP2-Orp1 expressing MN are affected by culture conditions, genotype, and degree of maturation. **A** Basal levels of H_2_O_2_ in the cytoplasm and mitochondria of the Cyto-roGFP2-Orp1 and Mito-roGFP2-Orp1 expressing MN measured on the differentiation day 29. **B** A basal level of H_2_O_2_ in the cytoplasm of the Cyto-roGFP2-Orp1 expressing MN measured on the differentiation day 20. **C** Basal levels of H_2_O_2_ in the cytoplasm and mitochondria of the Cyto-roGFP2-Orp1 and Mito-roGFP2-Orp1 expressing SOD1-G128R MN measured on the differentiation day 29 depending on the presence of neurotrophic factors in the culture medium during maturation. **D** Basal levels of H_2_O_2_ in the cytoplasm and mitochondria of the Cyto-roGFP2-Orp1 and Mito-roGFP2-Orp1 expressing MN in the B-27 deprived medium measured on the differentiation day 29. Data (N = 10-15, 18 for B-27 deprived samples) are the mean ± standard deviation. *** p < 0.001, **** p < 0.0001 with the Bonferroni correction for multiple comparisons where necessary; ### and #### denote p < 0.001 and p < 0.0001, respectively, in comparisons between the mitochondrial and cytoplasmic level of H_2_O_2_ of the same MN sample. The same datasets of the cytoplasmic and mitochondrial H_2_O_2_ levels in SOD1-G128R (-NTF) were used on the images **A** and **C**.

Standard culture conditions are not always beneficial for the redox studies because of the protective influence of the standard medium. We removed the B-27 supplement from the medium 24 h before live imaging of the MN and measured the cytoplasmic and mitochondrial H_2_O_2_ levels (Fig. 5d). We discovered that B-27 deprivation did not influence the cytoplasmic level of H_2_O_2_. Although MN cultured in the absence of B-27 demonstrated significant increase in the mitochondrial H_2_O_2_ level, which was, on average, 4 times higher than in the cytoplasm of the corresponding neurons. For the SOD1-G128R MN, the B-27 deprivation resulted in an elevation of both cytoplasmic and mitochondrial H_2_O_2_ levels to the values higher than in control K7-4Lf MN (Fig 5d).

### Antioxidants removal from the medium induces H_2_O_2_ accumulation in the cytoplasm of motor neurons

Removal of the antioxidants from the medium forces MN to rely on endogenous antioxidant systems for the ROS neutralization. We cultured motor neurons in the neuronal maintenance medium (supplemented with B-27 w/o antioxidants) for three days in addition to the differentiation protocol and performed live imaging of the MN (Fig. 6a-b). We observed an increase in the level of cytoplasmic H_2_O_2_ in the MN by 1.5-2.5 times. The mitochondrial H_2_O_2_ level did not change for control MN, iALS MN, and SOD1-D91A MN. However, together with increased cytoplasmic oxidation, SOD1-G128R MN demonstrated a rise in mitochondrial oxidation. This alteration in the H_2_O_2_ level was accompanied by visible changes in the SOD1- G128R MN morphology with the axon attrition and cytoplasmic vacuolization (Fig. 6c). Culturing of SOD1-G128R MN with the NTF reduced oxidation of the cytoplasm, but it did not manage to keep it to the normal level (Fig. 6d). The mitochondrial level of H_2_O_2_, again, was unaffected by the NTF addition.

**Fig. 6.**
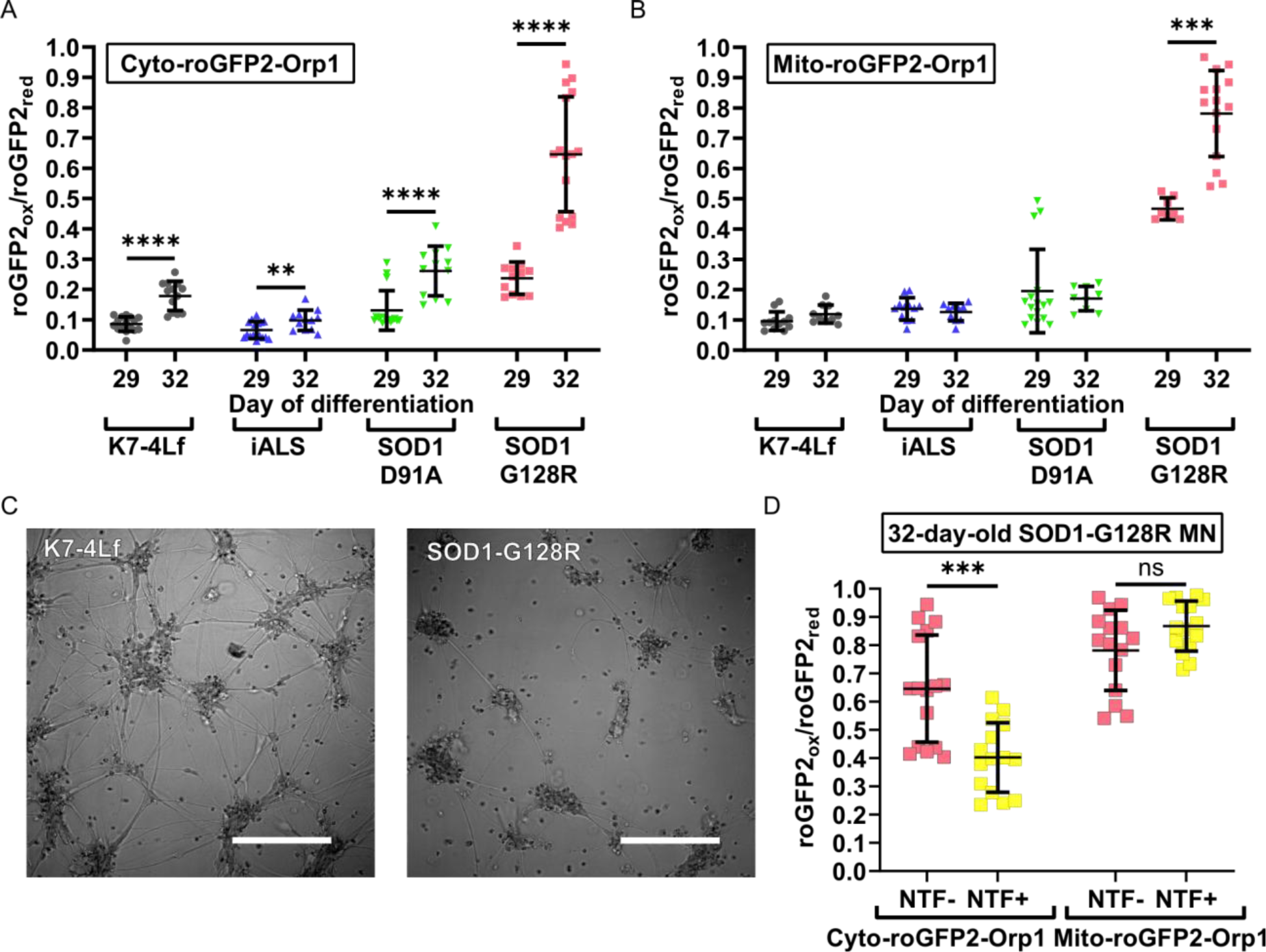
Effect of the antioxidant deprivation on the H_2_O_2_ level in the cytoplasm and mitochondria of motor neurons. A basal level of H_2_O_2_ in the cytoplasm (**A**) and mitochondria (**B**) of MN measured on the differentiation days 29 and 32. **C** Representative Images of the K7-4Lf and SOD1-G128R MN morphology in the antioxidant-deprived medium (differentiation day 32, DIC contrast), scale bar 200 μm. **D** Basal levels of H_2_O_2_ in the cytoplasm and mitochondria of SOD1-G128R MN measured on the differentiation day 32 depending on the presence of the neurotrophic factors in the culture medium. Data (N = 10-16) are the mean ± standard deviation. ns – non significant, ** p < 0.01,*** p < 0.001, **** p < 0.0001. The same datasets of cytoplasmic and mitochondrial H_2_O_2_ levels in SOD1-G128R (-NTF) were used on the images **A**, **B,** and **C**. The datasets used to present the cytoplasmic and mitochondrial oxidation on the differentiation day 29 for K7-4Lf, iALS, SOD1-D91A, and SOD1-G128R MN on the images **A** and **B** are the same as in Fig. 5A.

### Cyto-roGFP2-Orp1 biosensor expressed in motor neurons reflects the kinetics of H_2_O_2_ neutralization

H_2_O_2_ is a target molecule for the Cyto-roGFP2-Orp1 and Mito-roGFP2-Orp1 biosensors, and it directly affects the sensors’ properties and induces specific signals. At the same time, the addition of H_2_O_2_ to the cultured cells is a known method for stress induction [48]. The main advantage of the GE biosensors is that they allow measuring the H_2_O_2_ level in real-time, which is important for the study of dynamic cellular reactions. Thus, it is possible to record the process of H_2_O_2_ utilization by the cell. But the addition of deliberately lethal for the cells concentrations of H_2_O_2_ hinders the opportunity to compare the cellular response and its features caused by the genotype of interest. First, to find a maximum amount of hydrogen peroxide molecules that cultured MN can neutralize, staying alive, we tried a range of H_2_O_2_ concentrations. We found that MN was tolerant to 10 μM H_2_O_2_, but 25 μM H_2_O_2_ induced death of the cells (data not shown). Second, we tested whether the standard culture conditions distort the cellular response to the H_2_O_2_. We performed a live recording of the Cyto-roGFP2-Orp1-expressing MN reaction to the addition of 10 μM H_2_O_2_. These MN were either cultured in the standard NDM or starved in the B-27-deprived medium before the measurement (Fig. 7a). The overall reaction of the cells was similar regardless of the culture conditions; the MN expressing Cyto-roGFP2-Orp1 demonstrated oxidation followed by slow reduction, reflecting the change in the cytoplasmic H_2_O_2_ level. However, the reaction of MN cultured in the B-27-deprived medium before the experiment was more prominent. We detected a higher value of the maximum biosensor oxidation and faster reduction compared to the non-starved MN (Fig. b-c). Since the cellular reaction was affected by the components present in the standard medium, we conducted further measurements of the dynamic response on the cells that were starved before the experiment. Using the parameters of the experiment established earlier, we recorded the reaction of MN expressing the Mito-roGFP2-Orp1 biosensor to 10 μM H_2_O_2_ in real-time. We did not detect any changes in response to the exogenous H_2_O_2_; an oxidation value for the Mito-roGFP2-Orp1 biosensor remained constant during imaging, suggesting that the mitochondrial H_2_O_2_ level was also constant (Fig. 7d). The addition of H_2_O_2_ in higher concentrations (25 μM and 50 μM) induced mitochondrial oxidation but, as it was found earlier, damaged the neurons. This fact forced us to abandon the measurement of the dynamic response for the Mito-roGFP2-Orp1 sensor in further experiments (Fig. 7e).

**Fig. 7.**
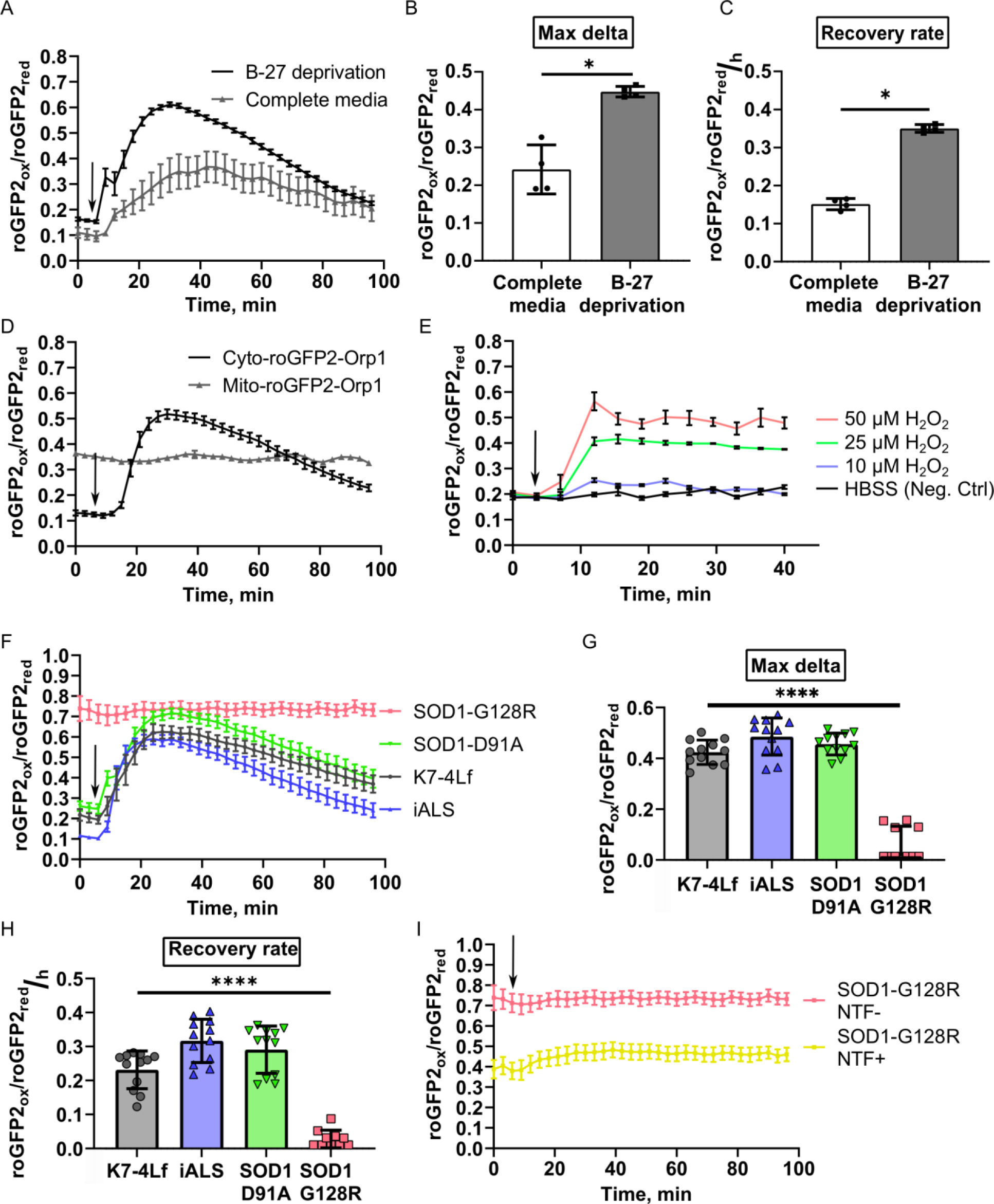
Kinetics of H_2_O_2_ utilization in live iPSC-derived motor neurons. **A** Response of the K7-4Lf MN cultured in the standard medium or B-27-deprived medium to the addition of 10 μM H_2_O_2_. Data (N 4) are the mean ± standard error of the mean (S.E.M.) Recovery rate (**B)** and maximum oxidation of the cytoplasm **(C)** of the K7-4Lf MN expressing Cyto-roGFP2-Orp1 to the H_2_O_2_ addition. Data (N 4) are the mean ± standard deviation (S.D.) **D** Response of the K7-4Lf MN expressing Cyto-roGFP2-Orp1 or Mito-roGFP2-Orp1 to the addition of 10 μM H_2_O_2_. Data (N 4) are the mean ± S.D. **E** Response of the K7-4Lf MN expressing Mito-roGFP2-Orp1 to the indicated amount of H_2_O_2_. Data (N 3) are the mean ± S.D. **F** Response of the K7-4Lf, iALS, SOD1-D91A, and SOD1- G128R MN expressing Cyto-roGFP2-Orp1 to the addition of 10 μM H_2_O_2_. Data (N 12) are the mean ± S.E.M. Recovery rate (**G**) and maximum oxidation of the cytoplasm (**H**) of the K7-4Lf, iALS, SOD1-D91A, and SOD1- G128R MN expressing Cyto-roGFP2-Orp1 after 10 μM H_2_O_2_ addition. Negative values in the SOD1-G128R sample were set to 0 on the graphs. Data (N 12) are the mean ± S. D. **I** Response of the SOD1-G128R MN expressing Cyto-roGFP2-Orp1 cultured in the standard medium or medium supplemented with neurotrophic factors to 10 μM H_2_O_2_. The moment of the H_2_O_2_ addition is marked by an arrow. Data (N 4) are the mean ± S.E.M. * p < 0.05, **** p < 0.0001. The same dataset was used in the images **F** and **I** for the representation of SOD1-G128R (-NTF) reaction to H_2_O_2_.

Next, we analyzed how the mutations introduced in *SOD1* affected neuronal reaction to the exogenous H_2_O_2_. We did not find any differences in the H_2_O_2_ utilization in the cytoplasm of Cyto-roGFP2-Orp1-expressing SOD1-D91A, iALS, and control (K7-4Lf) MN. The value of the maximum biosensor oxidation was similar between the cells, as well as the rate of recovery after the addition of H_2_O_2_ (Fig. 7f-h). Analysis of the response of the SOD1-G128R MN to the H_2_O_2_ revealed an aberrant reaction due to the initial high oxidation of the MN; the level of cytoplasmic oxidation remained constantly high before and after the addition of H_2_O_2_ (Fig. 7f, 7i). Although culturing of the SOD1-G128R MN with the NTF slightly reduced initial oxidation of the cytoplasm, it did not significantly affect the cell reaction to the H_2_O_2._ The Cyto-roGFP2-Orp1- expressing MN demonstrated moderate oxidation of the cytoplasm without signs of subsequent reduction (Fig. 7i).

### Motor neurons expressing the Cyto-roGFP2-Orp1 biosensor accumulate H_2_O_2_ in the cytoplasm due to glutamate-induced excitotoxicity

Glutamate excitotoxicity is one of the major mechanisms of the ALS development [49]. Excessive activation of the glutamate receptors leads to an increased Ca^2+^ influx, subsequent mitochondrial dysfunction, and apoptosis [50]. It is known that this process is accompanied by the increased ROS production, which connects it with the oxidative stress [51]. To test whether the Cyto-roGFP2-Orp1 and Mito-roGFP2-Orp1 biosensors can reflect redox imbalance caused by the excitotoxicity, we incubated the MN with monosodium glutamate and the glutamate reuptake inhibitor (PDC) for 5 days and measured the cytoplasmic and mitochondrial H_2_O_2_ levels. Since SOD1-G128R MN (both control and experimental) died shortly after the beginning of the experiment due to the reduced viability, the measurement was conducted only for K7-4Lf, SOD1-D91A, and iALS MN. We discovered that the glutamate treatment induced the accumulation of H_2_O_2_ in the cytoplasm, but not in the mitochondria, despite the known connection of mitochondrial dysfunction with the excitotoxicity (Fig. 8a-b). We observed mitochondrial oxidation in SOD1-D91A MN in both control and glutamate treated samples, although we were unable to determine if the oxidation was a hallmark of the SOD1 mutation or a technical artifact. The oxidation of the cytoplasm in K7-4Lf MN treated with glutamate was 28% higher, in iALS MN – by 41% higher, in SOD1-D91A MN – by 31.5% higher compared to the non-treated sample (Fig. 8a). We observed that iALS MN responded to the glutamate treatment more prominently than control MN. However, we did not find the same for SOD1-D91A (Fig. 8a), which makes us suggest that this effect was not caused by the *SOD1* mutation.

**Fig. 8.**
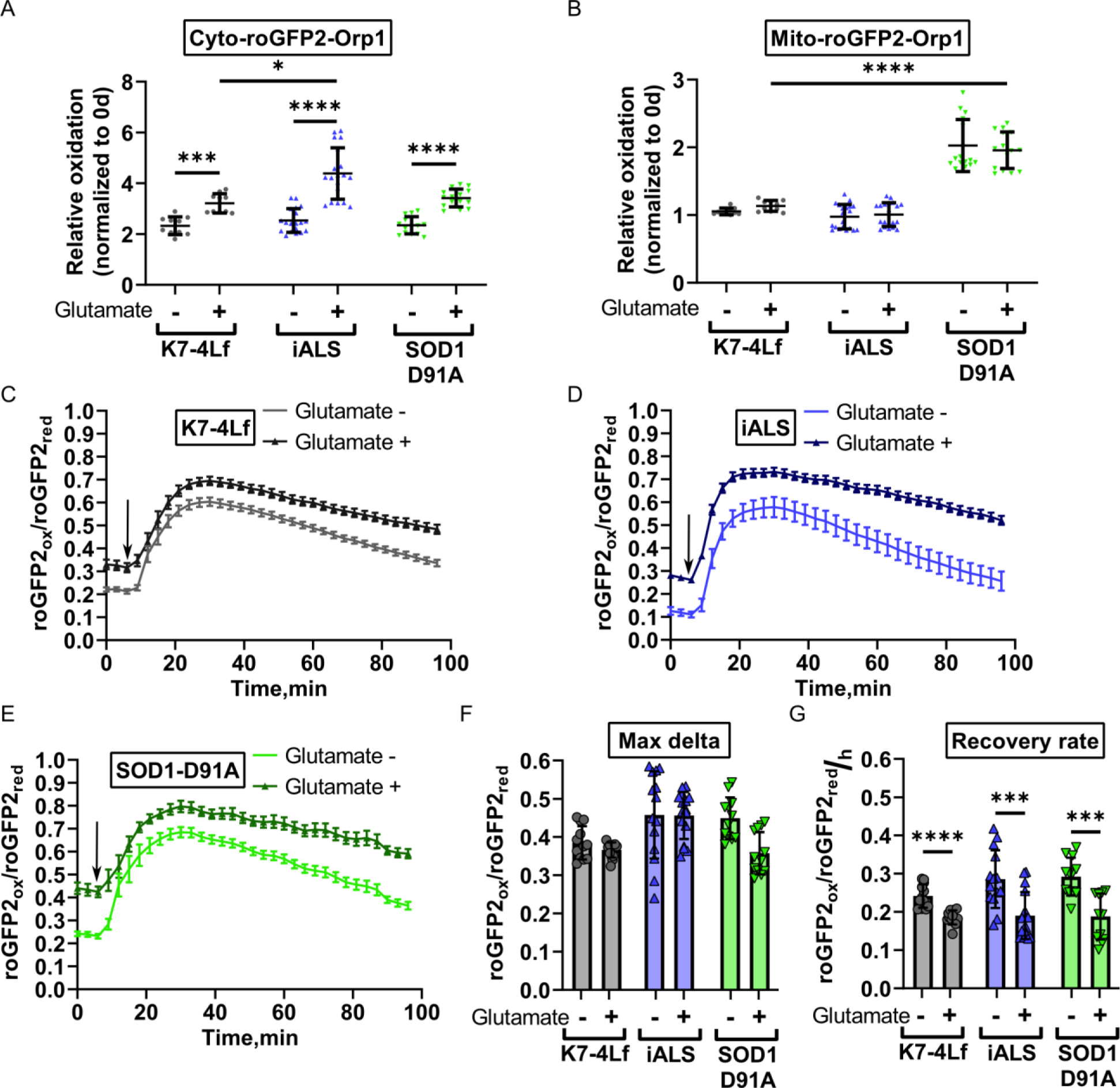
The reaction of the iPSC-derived motor neurons to the excitotoxicity. A basal level of H_2_O_2_ in the cytoplasm**A**) and mitochondria (**B**) of MN after 5-days incubation with glutamate. Data (N = 10-15) are normalized to the basal H_2_O_2_ level measured before the addition of glutamate and presented as the mean ± standard deviation (S.D.). The response of K7-4Lf (**C**), iALS (**D**), and SOD1-D91A (**E**) MN to the addition of 10 μM H_2_O_2_ (marked by an arrow). The reaction of MN cultured with glutamate is depicted in the darker graphs. Data (N 12) are the mean ± standard error of the mean (S.E.M.) Maximum oxidation of the cytoplasm (**F**) and recovery rate (**G**) of the K7-4Lf, iALS, and SOD1-D91A MN expressing Cyto-roGFP2-Orp1 after the addition of 10 μM H_2_O_2_. Data (N = 12) are the mean ± S. D. * p < 0.05, *** p < 0.001, **** p < 0.0001.

Further, we investigated how incubation with the monosodium glutamate affected dynamics of H_2_O_2_ utilization in the cytoplasm of the Cyto-roGFP2-Orp1-expressing MN. We found that MN treated with glutamate had reduced recovery rate after the H_2_O_2_ addition compared to the non- treated sample (Fig. 8c-g). We also observed the tendency towards the slower recovery of the mutant MN (36% slower for SOD1-D91A MN, 38% slower for iALS MN, 23% slower for K7- 4Lf MN), however, it was insignificant.

## DISCUSSION

In the present work, we established a valid approach for studying neurodegeneration in cell-based models using a combination of methods such as CRISPR/Cas9 genome editing, genetically encoded biosensors, and iPSC differentiation. Nowadays, GE biosensors are more often applied for research of different physiological and pathological processes [47,52,53] where they replaced molecular probes. In the case of redox processes, it provided information about dynamics of components of redox balance, e.g., hydrogen peroxide or GSH/GSSG ratio, in different cell compartments and tissues [23,47,54,55]. Since biosensors molecules are delivered inside the cells as plasmids for temporal expression or as viral vectors for sequence integration in the genome, the robustness of biosensors-related studies often relies on the efficiency of delivery and place of transgene integration. Here, we tried to improve the existing approach for biosensors experiments by target insertion of the cytoplasmic and mitochondrial H_2_O_2_ biosensors in combination with their controlled expression. We have generated a number of IPSC lines with the insertions of both the biosensor and transactivator essential for doxycycline-dependent expression, using the simultaneous delivery of CRISPR/Cas9 RNPs and donor plasmids.

We have discovered that a single copy of the biosensor is enough for producing a sufficient signal. Moreover, it allows achieving a similar dynamic range between samples, leading to more accurate data processing [56]. However, the mean dynamic range measured in the study for cytoplasmic and mitochondrial biosensors was comparable with the lowest numbers observed previously in other experiments [23, 47]. This indicates that the dynamic range is directly connected with the biosensor’s expression level, and more powerful promoters can improve this system [57]. The original design of the research suggested controllable induction of biosensors only on particular days during differentiation to avoid an excessive amount of fluorescent protein in the cell. However, we found a substantial decrease in fluorescence signal when it was induced only at the terminal stages of MN differentiation. Following measurement of mRNA expression confirmed a significant drop of the expression level of the biosensor, but not transactivator suggesting that maintaining the promoter in an active state prevents it from silencing [58]. Regular supplementation of doxycycline during differentiation helped to sustain a high level of the biosensors’ expression in MN. However, we still observed clones that partially lost their fluorescence and became mosaic, as well as clones with generally low fluorescence intensity (data not shown). All these suggest that chromatin remodeling occurring during differentiation affects transgenes expression in the *AAVS1* locus and sets a restriction on the type of transgenes that can be inserted into *AAVS1*. Although this problem can be overcome for biosensors [52], transgenes that are not supposed to be constantly active, such as cell-specific reporters, will not work well due to their promoter silencing [59, 60]. This fact underlines the necessity to discover new safe harbors in the human genome and test them not only in terms of safety but the ability to sustain a desirable level of transgene expression.

Using Cyto-roGFP2-Orp1 and Mito-roGFP2-Orp1 biosensors, we observed how MN genotype, culture conditions, and cell age affected the basal level of H_2_O_2_ in cytoplasm and mitochondria of the MN and their reaction to exogenous H_2_O_2_. We found that starvation, induced by the B-27 supplement removal from the medium, increased the mitochondrial level of H_2_O_2_ but generally did not affect the cytoplasmic level of H_2_O_2_ unless a severe genetic mutation was present (See further). Apparently, the rich composition of the standard NDM provides the environment that protects MN from various stressors coming from both outside and inside the culture system.

Starvation caused by B-27 deprivation induced moderate stress that served as a protection mechanism working through mitochondrial ROS as an activator of autophagy [61–63]. The lack of influence on the cytoplasmic level of H_2_O_2_ in normally functioning neurons suggests that this is a part of a natural response. Although the basal level of H_2_O_2_ in the cytoplasm was not affected by the starvation, MN reacted differently to the addition of H_2_O_2_ in the medium; starved MN demonstrated a higher amplitude of oxidation in response to H_2_O_2_ and a higher rate of recovery. B-27 deprivation made MN more susceptible to extra oxidation (thus, the higher amplitude of oxidation) due to the absence of protective antioxidants. At the same time, mobilization of the internal antioxidant system caused by starvation resulted in a more active reaction to the exogenous stress factors. It is unclear how long MN should be cultured in the B- 27 deprived medium before the stress becomes decompensated and whether it will help to distinguish MN with different genotypes. Some authors use deficit media on astrocyte-neuron cultures, and probably this approach can be applied for MN monocultures as well [35, 64].

Differences between Cyto-roGFP2-Orp1 and Mito-roGFP2-Orp1 reactions to starvation reflect relative independence of the mitochondrial and cytoplasmic antioxidant systems. Additionally, this fact is confirmed by the different reactions of Cyto-roGFP2-Orp1 and Mito-roGFP2-Orp1 to exogenous H_2_O_2_. The absence of significant changes in the signal of the Mito-roGFP2-Orp1 sensor and the presence of a normal reaction of the Cyto-roGFP2-Orp1 sensor in response to 10 μM H_2_O_2_ suggests that most exogenous H_2_O_2_ molecules are neutralized by the cytoplasmic antioxidant system and are not transferred in mitochondria. The reaction that only appears in response to lethal amounts of H_2_O_2_ suggests that exceeding of hydrogen peroxide level in the cytoplasm above the values that it can neutralize leads to a transfer of H_2_O_2_ molecules in mitochondria and apoptosis induction [50]. This effect has not been shown earlier, probably because previous studies did not focus on cell recovery and used large amounts of H_2_O_2_ for biosensor oxidation [23]. Possible insufficiently high dynamic range also can be the reason since it does not allow detecting small changes in H_2_O_2_ concentration. This also can explain the lack of mitochondria response to the glutamate: the power of the induced stress was not enough to provoke detectable changes in mitochondrial H_2_O_2_, but it didn’t mean that mitochondria were not affected. We believe that the application of more advanced versions of H_2_O_2_ biosensors with higher sensitivity and resolution in the future may be beneficial in such a study [65].

Correction of the mutation is more commonly used for to generate isogenic cell models [66, 67]. In this work, we did not aim to discover the effect of particular *SOD1* mutations but rather to test how the approaches used here can be applied for cell models. We tried to introduce the mutations in different parts of the *SOD1* gene, leading to the D91A and G128R substitutions in the polypeptide [40, 41]. And since no homozygous variants have been found, we selected the lines that have one allele with the target mutation and the other with large deletion (SOD1-D91A) or premature termination codon (SOD1-G128R) to maximize potential damage and make pathological phenotype more perceptible.

We found that MN derived from the SOD1-G128R iPSC line demonstrated significant differences in functioning that were detected by the biosensors. The most apparent is the higher level of cytoplasmic and mitochondrial oxidation that was found in the MN. Moreover, SOD1- G128R MN reacted more prominent than others to the stress tests such as starvation, induced by B-27 deprivation and culturing in the antioxidant-deprived medium. Notably, immature SOD1- G128R MN did not show differences in cytoplasmic H_2_O_2_, suggesting that the pathological phenotype requires time to develop, as it happens in ALS patients, who usually develop the symptoms after a certain age [68]. Neurotrophic factors affected cytoplasmic but not mitochondrial H_2_O_2_ level, although it became less apparent in the antioxidant-deprived medium. This suggests that SOD1-G128R MN functioning only relies on a highly protective culture system, and these MN are unable to sustain their viability without additional help. An elevated level of mitochondrial H_2_O_2_ regardless of the NTF addition shows the presence of unresolved mitochondrial malfunction that could not be corrected at that point. Presumably, mitochondrial dysfunction of SOD1-G128R MN resulting from the presence of mutant *SOD1* in the cell leads to the redox imbalance and H_2_O_2_ accumulation inside the mitochondria at first [69]. Then, when the H_2_O_2_ level reaches the limit of buffering capacity in mitochondria, H_2_O_2_ enters the cytoplasm, probably due to the mitochondria death and fragmentation, initiating apoptosis. The presence of mitochondrial dysfunction also explains the slower axon growth since the lack of energy due to the dysfunction forces the cell to focus on survival rather than rebuilding the cytoskeleton [70].

Despite the significant, albeit not so prominent, decrease in axonal growth rate observed for SOD1-D91A and iALS MNs, cytoplasmic and mitochondrial H_2_O_2_ levels did not demonstrate significant differences associated with *SOD1* mutation from the control. Patient-specific iALS MN demonstrated even lower H_2_O_2_ levels in some measurements, but the data was inconsistent and did not reproduce in SOD1-D91A MN, suggesting that this effect was related to the line specificity since iALS has a different origin. This data highlights the necessity to apply isogenic iPSC since such differences can lead to wrong conclusions. Glutamate-induced excitotoxicity intended to uncover mutant MN malfunction did not help as well. Although glutamate treated neurons demonstrated accumulation of H_2_O_2_ in the cytoplasm and slower recovery rate after H_2_O_2_ addition, no differences between mutant and control MN were observed, suggesting that the power of stress or the degree of MN maturation were insufficient for the pathological phenotype to reveal. Nevertheless, different degree of disturbances observed in SOD1-D91A and SOD1-G128R motor neurons in the study allows concluding that these neurons recapitulate the clinical effect of such mutations in terms of different levels of severity of the symptoms.

## CONCLUSION

We developed a cell-based platform that allowed us to study redox balance in the live motor neurons with the detection of basal oxidation of the cytoplasm and mitochondria and the dynamic response of the cells to the stressors. Using isogenic iPSCs, we confirmed that mutations affecting different parts of the *SOD1* sequence have a different impact on motor neurons state *in vitro* as it has been observed in patients. Our work presents a new approach for the application of cell-based disease models for research and can be used to generate other similar models. We expect this approach to be expanded to include other disease-associated mutations or the biosensors of other cellular processes.

## ABBREVIATIONS

AAVS1 – adeno-associated virus site 1; ALS – amyotroptic lateral sclerosis; ARMS – amplification-refractory mutation system; CRISPR/Cas9 – clustered regularly interspaced short palindromic repeats/ CRISPR-associated; DTT – dithiothreitol; GE biosensors – genetically encoded biosensors; gRNA – guide RNA; HBSS – Hank’s balanced salt solution; iPSC – induced pluripotent stem cells; MN – motor neurons; NDefM – neuronal deficit medium; NDM – neuronal differentiation medium; NTF – neurotrophic factors; RNP – ribonucleoprotein; roGFP2 – reduction-oxidation sensitive green fluorescent protein 2; ROS – reactive oxygen species; rtTA – reverse tetracycline-controlled transactivator; SOD1 – superoxide dismutase 1; ssODN - single-stranded oligodeoxynucleotide.

## SUPPLEMENTARY INFORMATION

**Additional file 1: Table S1**. List of iPSC lines used in the study.

**Additional file 2: Table S2**. List of primers and oligonucleotides used in the study.

**Additional file 3: Table S3**. List of antibodies used in the study.

**Additional file 4: Supplementary Figure 1**. Graphical description of live MN sample preparation for microscopy. **A** Schematic representation of the protocol for MN sample preparation. **B** 3D reconstruction of the cell layer (differentiation day 29) obtained with the protocol. Morphology of MN seeded for the maturation on the top of the Matrigel (**C**) or inside layer of 33% Matrigel (**D**) at the indicated days of differentiation.

**Additional file 5: Supplementary Figure 2**. SOD1-D91A and SOD1-G128R iPSC lines characterization. **A** Pluripotent features of the SOD1-D91A and SOD1-G128R iPSC lines. Immunofluorescent staining for transcriptional factors: SOX2, OCT4, NANOG, and surface antigen TRA-1-60. Nuclei are visualized with DAPI. Scale bar 100 μM. IPSC colony morphology (TL – transmitted light), scale bar 150 μM. Alkaline phosphatase (AP) staining, scale bar 100 μM. **B** RT-qPCR analysis of mRNA expression of *SOX2*, *OCT4,* and *NANOG* in SOD1-D91A and SOD1-G128R iPSCs. Data (N = 3) are normalized to the parental iPSC line (K7-4Lf) and presented as the mean ± standard deviation. **C** Immunofluorescent staining of the products of *in vitro* spontaneous differentiation for endodermal (CK18), ectodermal (NF200), and mesodermal (ɑSMA) markers. Nuclei are visualized with DAPI. Scale bar 100 μM. **D** Partial sequence of exon 4 of the *SOD1* gene of SOD1-D91A with the corresponding amino acid sequence. WT ref – reference sequence (parental iPSC line K7-4Lf); D91A allele– allele with the c.272A>C single nucleotide substitution; Del allele – allele with 105-nucleotide deletion. **E**. Partial sequence of exon 5 of the *SOD1* gene of SOD1-G128R with the corresponding amino acid sequences. WT ref. – reference sequence (parental iPSC line K7-4Lf); G128R allele – allele with the c.382G>C single nucleotide substitution; Stop allele – allele with premature termination codon. **F** Mycoplasma contamination detection with specific primers. Neg. ctrl – H_2_O, Pos. ctrl – mycoplasma contaminated cell line. Karyotyping of the SOD1-D91A (**G**) iPSC line and SOD1- G128R (**H**) iPSC line.

**Additional file 6: Supplementary Figure 3**. pCyto-roGFP2-Orp1-donor (**A**) and pMito- roGFP2-Orp1-donor (**B**) plasmid maps. Schematic maps of the donor plasmids used for CRISPR/Cas9 target insertion of the biosensors in the *AAVS1* locus. Basic elements: homologous arms, puromycine resistance gene, promoters, Tet-On elements for doxycycline-controllable expression, and biosensors’ functional elements are present. All maps were made with SnapGene®.

**Additional file 7: Supplementary Figure 4** SOX2 and SSEA4 staining of the transgenic iPSC clones expressing Cyto-roGFP2-Orp1 (_Cyto) or Mito-roGFP2-Orp1 (_Mito) biosensors. Nuclei are visualized with DAPI, scale bar 100 μM.

**Additional file 8: Supplementary Figure 5** Immunofluorescent staining of iPSC-derived motor neurons (differentiation day 28) for ChAT, MNX1, and ISL1. Nuclei are visualized with DAPI. Scale bar 200 μM.

## ACKNOWLEDGEMENTS

Authors wish to thank the Microscopy Center of Biological Objects of the Siberian Branch of the Russian Academy of Sciences for granting access to microscopic equipment and SB RAS Genomics Core Facility for Sanger sequencing of the samples.

## AUTHORS’ CONTRIBUTIONS

EIU and SPM designed the research. EIU performed the experiments and analyzed the data. SPM and SMZ obtained the funding. EIU drafted the manuscript and SPM and SMZ revised and approved it. All authors have read and agreed to the published version of the manuscript.

## AVAILABILITY OF DATA AND MATERIALS

The datasets used and/or analyzed during the current study are available from the corresponding authors on reasonable request.

## ETHICS APPROVAL AND CONSENT TO PARTICIPATE

The work regarding genetic modification of the iPSC lines, generated from the patients’ materials has been approved by the Research Ethics Committee of FSBI Federal Neurosurgical Center (Novosibirsk, Russia), protocol number 1 14/03/2017.

## CONSENT FOR PUBLICATION

Not applicable.

