## Supplementary material for "A cell-based platform for oxidative stress monitoring in motor neurons using genetically encoded biosensors of H_2_O_2_": Supplementary table 1. List of iPSCs used in the study.docx

**Table S1.** List of iPSC lines used in the study

| Name | Genotype | Generation and characteristics | HpscReg # | Remarks |
| --- | --- | --- | --- | --- |
| K7-4Lf | *SOD1^WT/WT^* | (Malakhova et al., 2020) | ICGi022-A | Exome sequencing data:  https://www.ncbi.nlm.nih.gov/sra/?term=SRR11413027; isogenic control to the ICGi022-A-1 and ICGi022-A-2 |
| iALS | *SOD1^D91A/D91A^* | (Ustyantseva et al., 2020) | ICGi04-A |  |
| SOD1-D91A | *SOD1^D91A/Δ105^* | Present study | ICGi022-A-1 | Genetically modified subclone of ICGi-022-A. Both alleles of the *SOD1* gene were modified by CRISPR/Cas9 with [c.272A>C] + [c.238+675_281del] mutations. |
| SOD1-G128R | *SOD1^G128R/K129X^* | Present study | ICGi022-A-2 | Genetically modified subclone of ICGi022-A. Both alleles of the *SOD1* gene were modified by CRISPR/Cas9 with [c.382G>C] + [c.385_386 delinsTG] mutations, leading to the G128R and K129* substitutions in the protein sequence, respectively. |
| K7-4Lf-Cyto1,  K7-4Lf-Cyto2,  K7-4Lf-Cyto3 | *SOD1^WT/WT^* | Present study | - | Contain transgenes of the cytosolic H_2_O_2_ biosensor Cyto-roGFP2-Orp1 and tetracycline-transactivator (rtTA) in *AAVS1* locus |
| K7-4Lf-Mito1, K7-4Lf-Mito2, K7-4Lf-Mito3 | *SOD1^WT/WT^* | Present study | - | Contain transgenes of the mitochondrial H_2_O_2_ biosensor Mito-roGFP2-Orp1 and tetracycline-transactivator (rtTA) in *AAVS1* locus |
| iALS-Cyto1,  iALS-Cyto2, iALS-Cyto3 | *SOD1^D91A/D91A^* | Present study | - | Contain transgenes of the cytosolic H_2_O_2_ biosensor Cyto-roGFP2-Orp1 and tetracycline-transactivator (rtTA) in *AAVS1* locus |
| iALS-Mito1, iALS-Mito2, iALS-Mito3 | *SOD1^D91A/D91A^* | Present study | - | Contain transgenes of the mitochondrial H_2_O_2_ biosensor Mito-roGFP2-Orp1 and tetracycline-transactivator (rtTA) in *AAVS1* locus |
| SOD1-D91A-Cyto1,  SOD1-D91A-Cyto2, SOD1-D91A-Cyto3 | *SOD1^D91A/Δ105^* | Present study | - | Contain transgenes of the cytosolic H_2_O_2_ biosensor Cyto-roGFP2-Orp1 and tetracycline-transactivator (rtTA) in *AAVS1* locus |
| SOD1-D91A-Mito1, SOD1-D91A-Mito2, SOD1-D91A-Mito3 | *SOD1^D91A/Δ105^* | Present study | - | Contain transgenes of the mitochondrial H_2_O_2_ biosensor Mito-roGFP2-Orp1 and tetracycline-transactivator (rtTA) in *AAVS1* locus |
| SOD1-G128R-Cyto1, SOD1-G128R-Cyto2, SOD1-G128R-Cyto3 | *SOD1^G128R/K129X^* | Present study | - | Contain transgenes of the cytosolic H_2_O_2_ biosensor Cyto-roGFP2-Orp1 and tetracycline-transactivator (rtTA) in *AAVS1* locus |
| SOD1-G128R-Mito1, SOD1-G128R-Mito2, SOD1-G128R-Mito3 | *SOD1^G128R/K129X^* | Present study | - | Contain transgenes of the mitochondrial H_2_O_2_ biosensor Mito-roGFP2-Orp1 and tetracycline-transactivator (rtTA) in *AAVS1* locus |
