## Supplementary material for "A cell-based platform for oxidative stress monitoring in motor neurons using genetically encoded biosensors of H_2_O_2_": Supplementary table 2. List of primers and oligos used in the study.docx

**Table S2.** List of the primers and oligonucleotides used in the study

| Name | Sequence, 5’-3’ | Target |
| --- | --- | --- |
| Oligonucleotides | | |
| AAVS1 crisprRNA | GUCACCAAUCCUGUCCCUAGGUU  UUAGAGCUAUHCU | crisprRNA targeting *AAVS1* |
| SOD1-4 crisprRNA | GACUGCUGACAAAGAUGGUGGUU  UUAGAGCUAUGCU | crisprRNA targeting exon 4 of *SOD1* (c.272A>C substitution) |
| SOD1-5 crisprRNA | GCAGAUGACUUGGGCAAAGGGUU  UUAGAGCUAUGCU | crisprRNA targeting exon 5of *SOD1* (c.382G>C substitution) |
| D91A-ssODN donor | A*G*A*TCACAGAATCTTCAATAGAC  ACGTCGGCCACACCATCTTTGGCAG  CAGTCACATTGCCCAAGTCTCCAAC  ATGCCTAA TAATGAA*A*A*A | ssODN donor sequence (c.272A>C substitution in *SOD1*) * - Phosphorothioate bonds |
| G128R-ssODN donor | G*T*T*TCCTGTCTTTGTACTTTCTTCAT  TTCCACCTTTGCGCAAGTCATCTGC  TTTTTCATGAACCTGTAAAAAATTT  TAAGAAGATAAC*T*T*T | ssODN donor sequence (c.382G>C substitution in *SOD1*) * - Phosphorothioate bonds |
| DNA-probes for c.382G>C detection | | |
| G128G-WT-FAM | FAM-CCTTTGCCCAAGTCATCTGC-BHQ1 | Wild type allele detection |
| G128R-mut-VIC | VIC-CCTTTGCGCAAGTCATCTGC-BHQ2 | c.382G>C mutant allele detection |
| Primers | | |
| SOD1-D91A-F | ATATCAGAGGCCTTGGGACATAG | Amplification of the exon 4 of *SOD1* |
| SOD1-D91A-R | TGAACTGCAAGTACAGTTTATCTGG |  |
| SOD1-G128R-F | TGTCTTTGCAACACCAAGAAA | Amplification of the exon 5 of *SOD1* |
| SOD1-G128R-R | TTCACAGGCTTGAATGACAAA |  |
| B2M-F | TAGCTGTGCTCGCGCTACT | Housekeeping reference gene for qPCR |
| B2M-R | TCTCTGCTGGATGACGTGAG |  |
| RPL13-F | CCTGGAGGAGAAGAGGAAAGAGA | Housekeeping reference gene for qPCR |
| RPL13-R | TTGAGGACCTCTGTGTATTTGTCAA |  |
| GAPDH-F | TGTTGCCATCAATGACCCCTT | Housekeeping reference gene for qPCR |
| GAPDH-R | CTCCACGACGTACTCAGCG |  |
| HPRT1-F | GACTTTGCTTTCCTTGGTCAGG | Housekeeping reference gene for qPCR |
| HPRT1-R | AGTCTGGCTTATATCCAACACTTCG |  |
| OCT4-F | CTTCTGCTTCAGGAGCTTGG | Pluripotency markers expression |
| OCT4-R | GAAGGAGAAGCTGGAGCAAA |  |
| SOX2-F | GCTTAGCCTCGTCGATGAAC |  |
| SOX2-R | AACCCCAAGATGCACAACTC |  |
| NANOG-F | CAGCCCCGATTCTTCCACCAGTCCC |  |
| NANOG-R | CGGAAGATTCCCAGTCGGGTTCACC |  |
| HB9-F | GTCCACCGCGGGCATGATCC | Motor neuron markers expression |
| HB9-R | TCTTCACCTGGGTCTCGGTGAGC |  |
| CHAT-F | GGAGGCGTGGAGCTCAGCGACACC |  |
| CHAT-R | CGGGGAGCTCGCTGACGCAGTCTG |  |
| ISL-F | AGCAGCCCAATGACAAAACT |  |
| ISL-R | CTGAAAAATTGACCAGTTGCTG |  |
| Myco-F | GGGAGCAAACAGGATTAGATACCCT | Mycoplasma contamination detection |
| Myco-R | TGCACCATCTGTCACTCTGTTAACCTC |  |
| SOD1- qPCR-G128R-F | GTAGTGATTACTTGACAGCC | Amplification of the part of the exon 5 of SOD1 for c.382G>C detection |
| SOD1- qPCR-G128R-R | CAATTACACCACAAGCCAA |  |
| roGFP2-qPCR-F | TAGACGTTGTGGCAGTTGTAG | Biosensors’ expression |
| roGFP2-qPCR-R | GCTGAAGGGCATCGACTT |  |
| rtTA-qPCR-F | GGACAGGCATCATACCCACTT | Transactivator expression |
| rtTA-qPCR-R | AGAGCACAGCGGAATGACTT |  |
| HA_L-OUT | CCGGACCACTTTGAGCTCTAC | Target insertion of the biosensor/transactivator in *AAVS1* |
| Neo_in-R | GCCCAGTCATAGCCGAATAG | + HA_L_OUT, Target insertion of the transactivator |
| Puro_in-R | AGGCGCACCGTGGGCTTGTAC | + HA_L_OUT, Target insertion of the biosensor |
| M13-R | CAGGAAACAGCTATGAC | +Neo_in-R, off-target insertions of the AAVS1-Neo-M2rtTA donor |
| M13-F | GTAAAACGACGGCCAGT | +Puro_in-R, off-target insertions of the Cyto-roGFP2-Orp1/ Cyto-roGFP2-Orp1 donor |
| D91A-inner-F | CTTGGGCAATGTGACTGCCGA | c.272A>C detection in *SOD1* using Tetra-primer ARMS-PCR |
| D91A-inner-R | ATCGGCCACACCATCTTCGG |  |
| D91A-outer-F | TGATGTTTAGTGGCATCAGCCCTAATC |  |
| D91A-outer-R | GAAACCGCGACTAACAATCAAAGTG  AA |  |
