## Supplementary material for "A cell-based platform for oxidative stress monitoring in motor neurons using genetically encoded biosensors of H_2_O_2_": Supplementary table 3 List of antibodies used in the study.docx

**Table S3.** List of antibodies used in the study

| Antibody | Vendor | Cat # | Host | Dilution |
| --- | --- | --- | --- | --- |
| **Primary antibody** | | | | |
| Pluripotency markers | | | | |
| OCT4 | BD Transduction Lab | 611202 | Mouse monoclonal | 1:200 |
| NANOG | Abcam | ab62734 | Rabbit polyclonal | 1:200 |
| SOX2 | Cell Signaling | 3579 | Rabbit polyclonal | 1:400 |
| SSEA4 | Abcam | ab16287 | Mouse monoclonal | 1:50 |
| TRA-1-60 | Abcam | ab16288 | Mouse monoclonal | 1:200 |
| Ecto-, meso-, endoderm markers | | | | |
| NF200 | Sigma | N4142 | Rabbit polyclonal | 1:500 |
| aSMA | Dako |  | Mouse monoclonal IgG2a |  |
| CK18 | EMD Millipore | MAB3236 | Mouse monoclonal IgG1 | 1:200 |
| Motor neuron markers | | | | |
| TUJ1 | Biolegend | 801213 | Mouse monoclonal IgG2 | 1:1000 |
| HB9 | EMD Millipore | ABN174 | Rabbit polyclonal | 1:1000 |
| ISL1 | Abcam | ab109517 | Rabbit polyclonal | 1:500 (1:100 flow cytometry) |
| ChAT | EMD Millipore | AB144P | Rabbit polyclonal | 1:100 |
| **Secondary antibody** | | | | |
| Alexa Fluor 488 goat anti rabbit IgG (H+L) | Thermo Fisher Scientific | A11008 | Goat polyclonal | 1:400 |
| Alexa Fluor 568 goat anti rabbit IgG (H+L) | Thermo Fisher Scientific | A11011 | Goat polyclonal | 1:400 (1:2000 flow cytometry) |
| Alexa Fluor 488 goat anti mouse IgG (H+L) | Thermo Fisher Scientific | A11029 | Goat polyclonal | 1:400 |
| Alexa Fluor 568 goat anti mouse IgG (H+L) | Thermo Fisher Scientific | A11004 | Goat polyclonal | 1:400 |
| Alexa Fluor 568 goat anti mouse IgM | Thermo Fisher Scientific | A21043 | Goat polyclonal | 1:400 |
| Alexa Fluor 488 rabbit anti goat IgG (H+L) | Thermo Fisher Scientific | A11078 | Goat polyclonal | 1:400 |
