## Supplementary figures and images for "A cell-based platform for oxidative stress monitoring in motor neurons using genetically encoded biosensors of H_2_O_2_"

### Supplementary fig 1.3D matrix graphical description.png

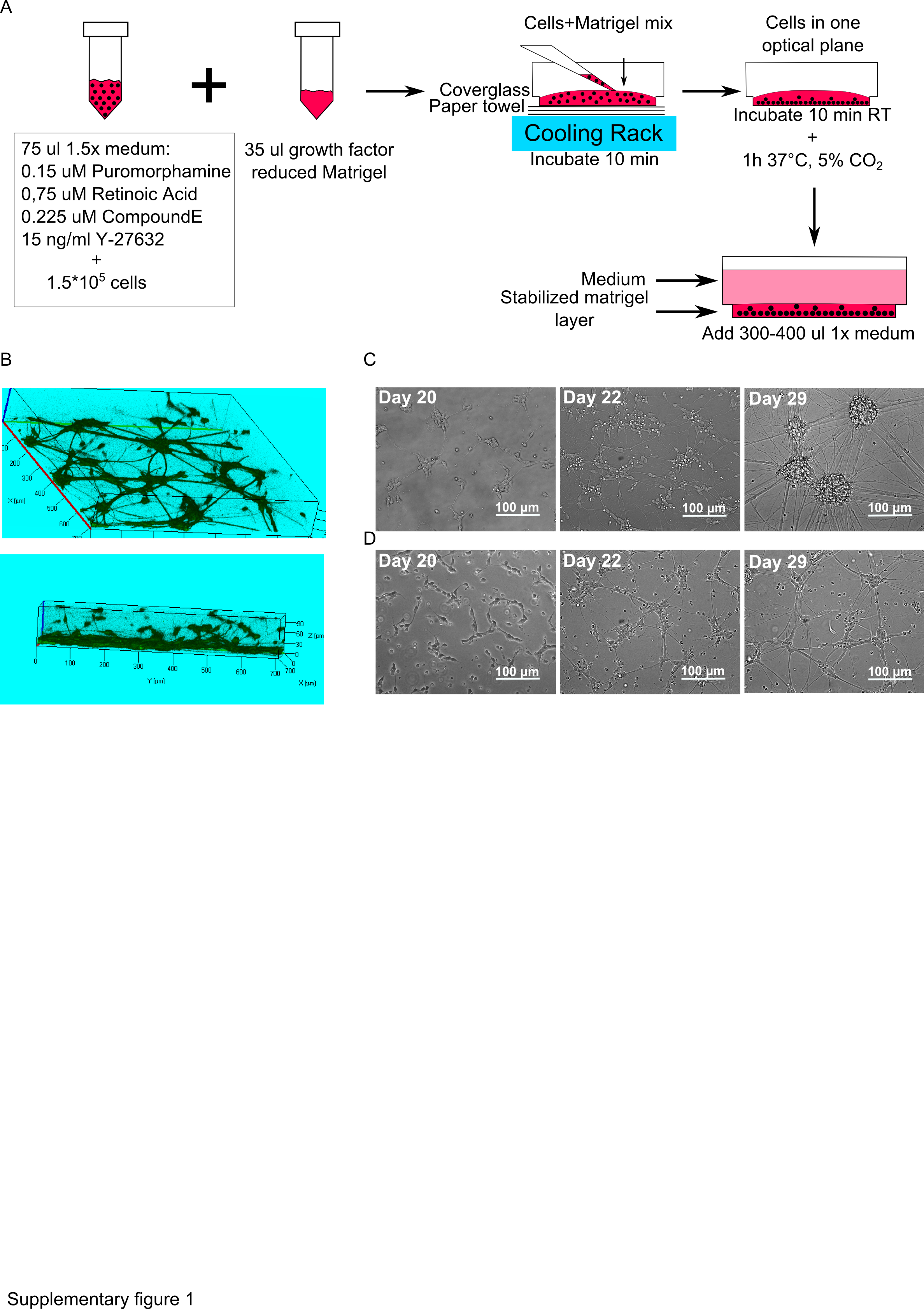

### Supplementary fig 2. Characterisation of the SOD1 mutant iPSCs.png

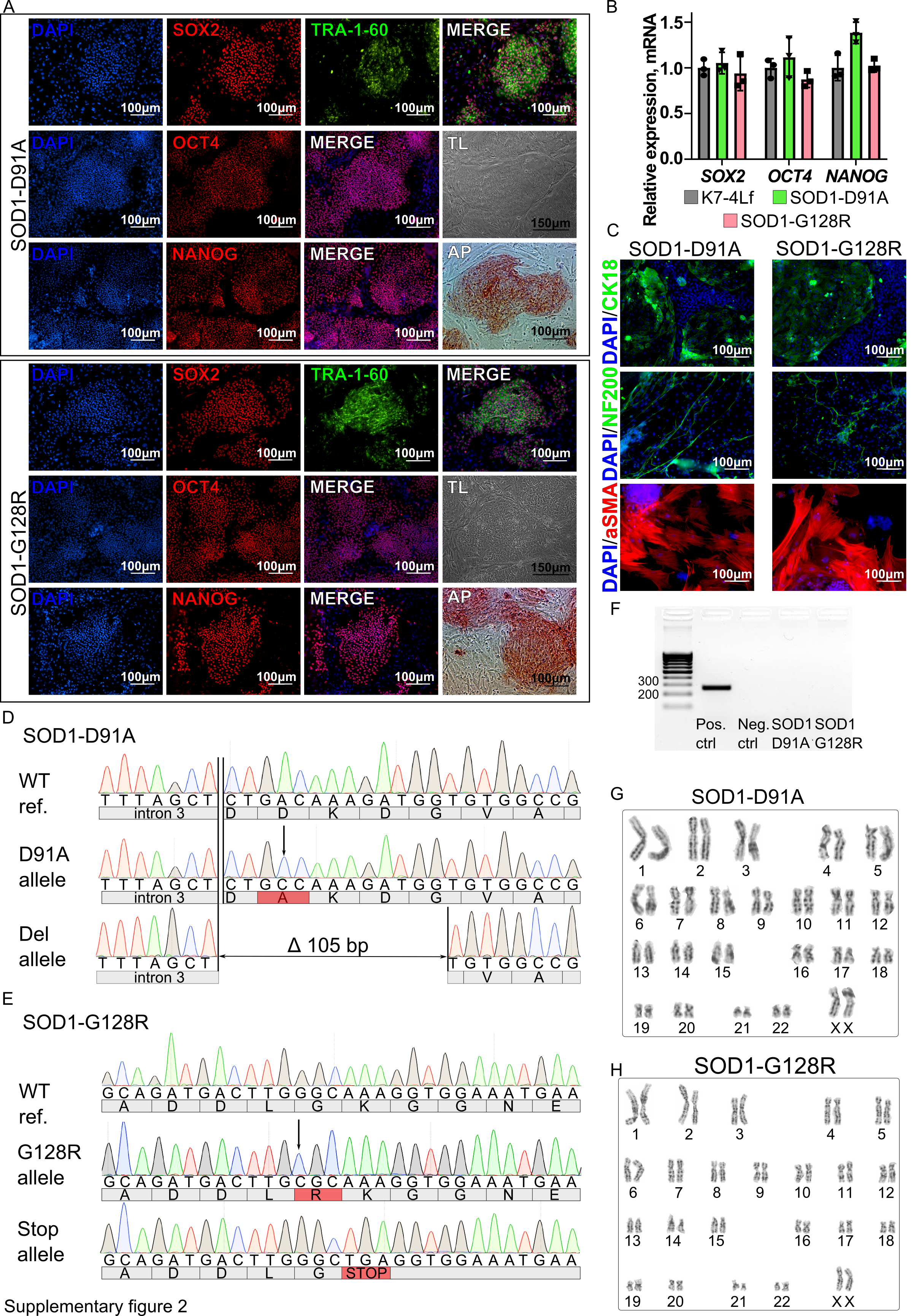

### Supplementary fig 3. Donor plasmid maps.png

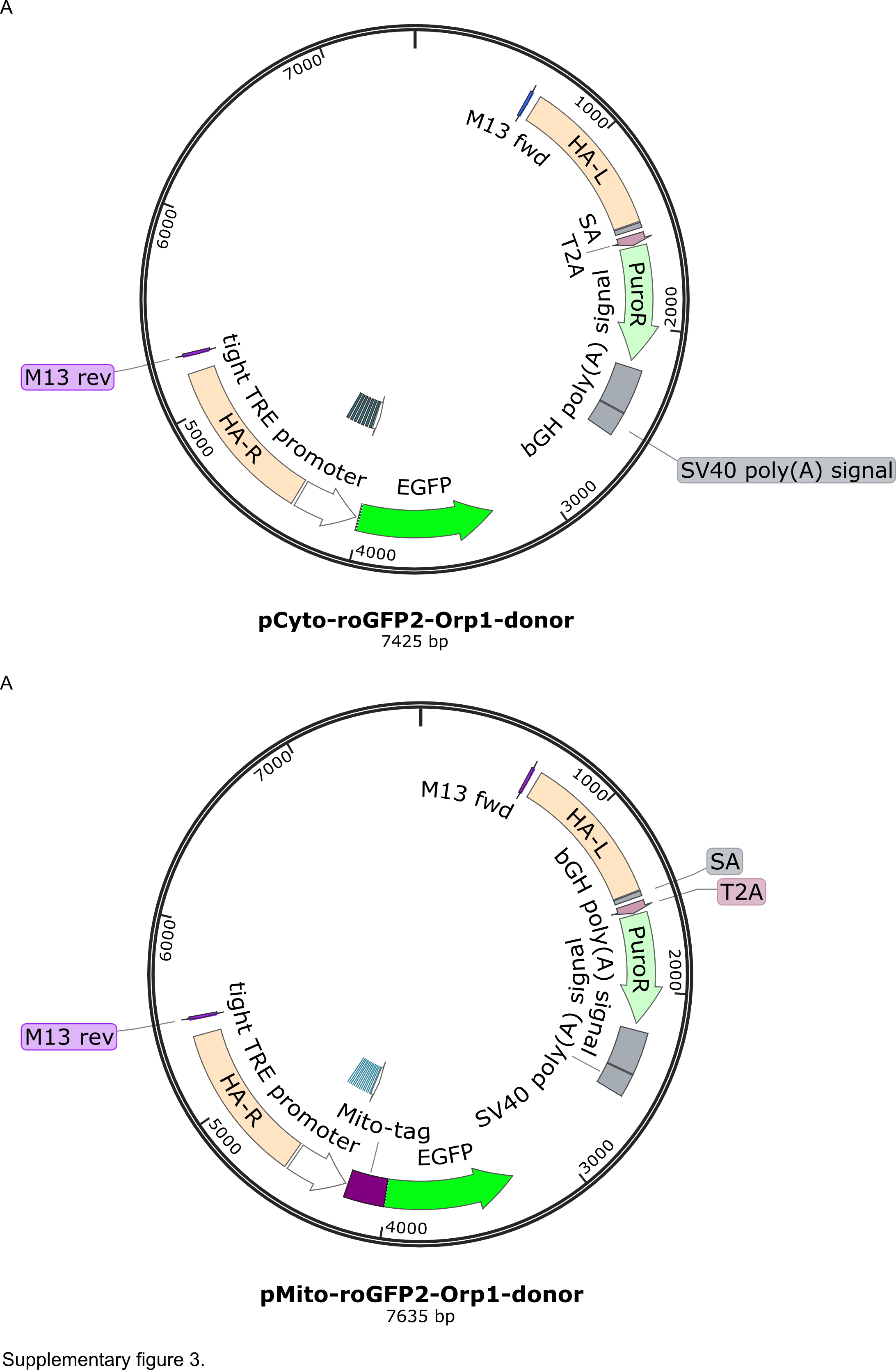

### Supplementary fig 4. SOX1 and SSEA4 staining of the biosensors' expressin iPSCs.png

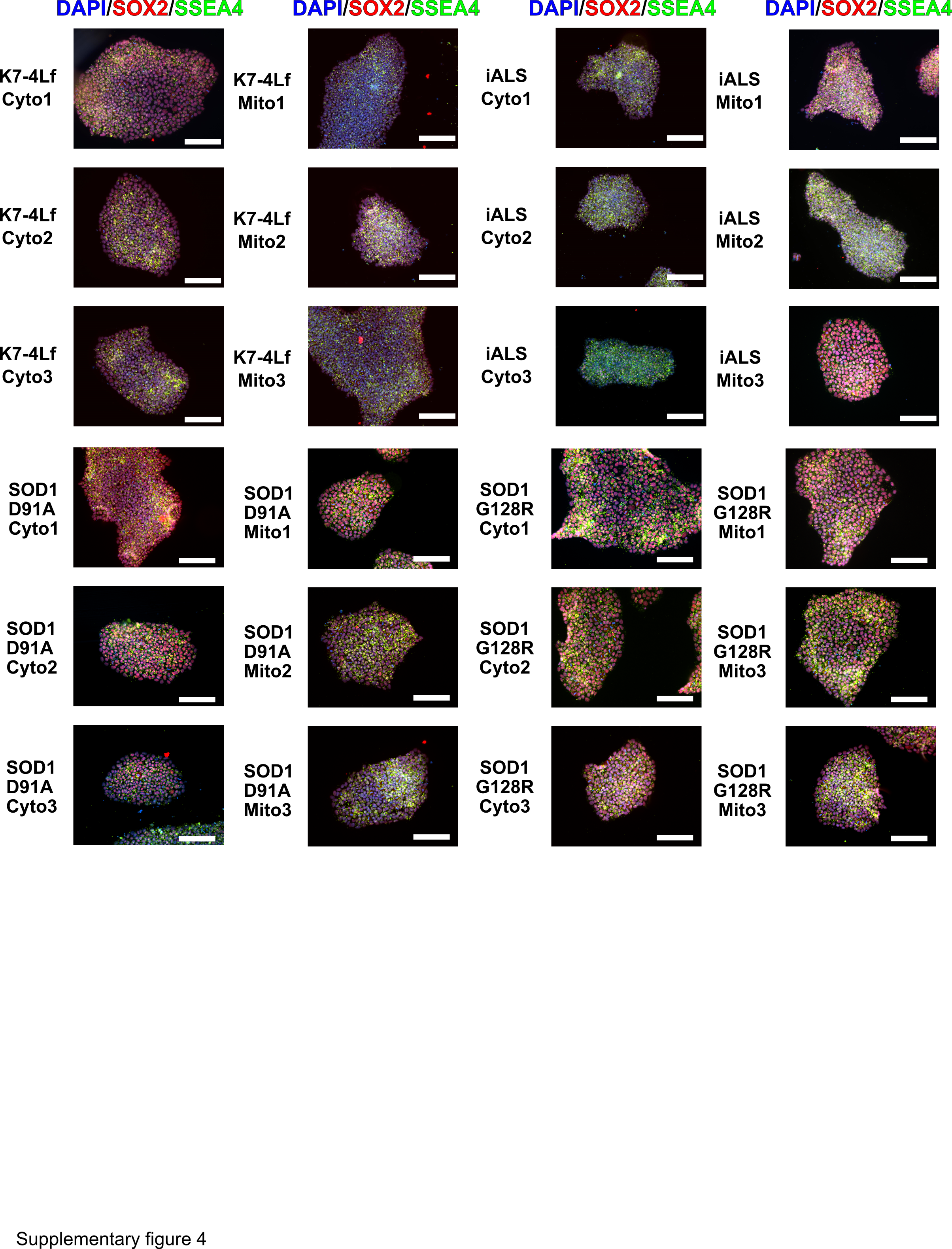

### Supplementary fig 5. MN markers expanded.png

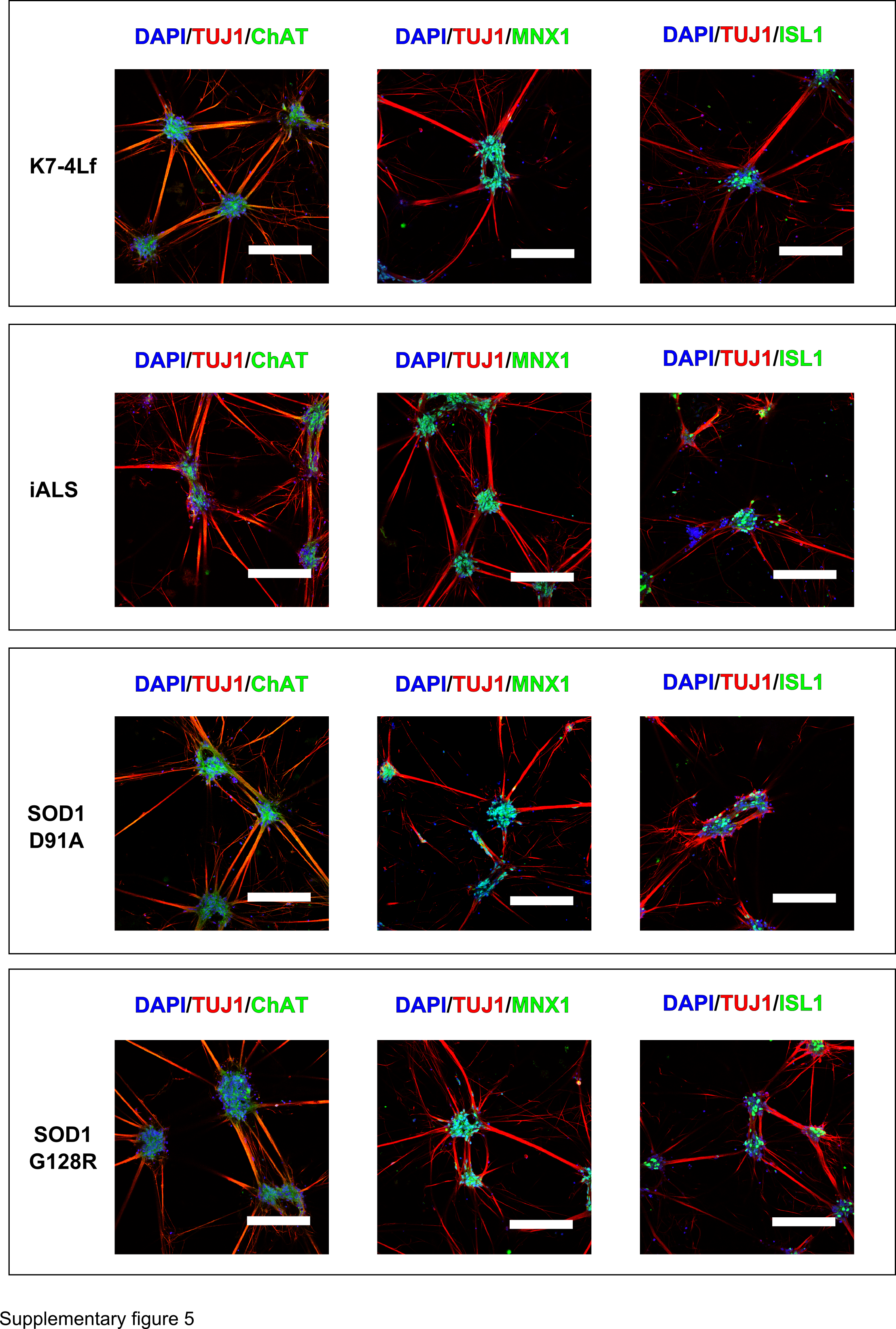
